# Innate immune system signaling and CD11b^+^CD11c^+^CD103^+^ cell migration to the brain underlie changes in mouse behavior after microbial colonization

**DOI:** 10.1101/2024.07.03.600853

**Authors:** Vivek Philip, Narjis Kraimi, Hailong Zhang, Jun Lu, Giada De Palma, Chiko Shimbori, Kathy D. McCoy, Siegfried Hapfelmeier, Olivier P. Schären, Andrew J Macpherson, Fernando Chirdo, Michael G. Surette, Elena F. Verdu, Fang Liu, Stephen M Collins, Premysl Bercik

## Abstract

**Background and Aims:** Accumulating evidence suggests the microbiota is a key factor in disorders of gut-brain interaction (DGBI), by affecting host immune and neural systems. However, the underlying mechanisms remain elusive due to their complexity and clinical heterogeneity of patients with DGBIs. We aimed to identify neuroimmune pathways that are critical in microbiota-gut-brain communication during de novo gut colonization.

**Methods:** We employed a combination of gnotobiotic and state-of-the-art microbial tools, behavioral analysis, immune and pharmacological approaches. Germ-free wild type, MyD88^−/−^ Ticam1^−/−^ and SCID mice were studied before and after colonization with specific pathogen-free microbiota, Altered Schaedler Flora, E. coli or S. typhimurium (permanent or transient colonizers). TLR agonists and antagonists, CCR7 antagonist or immunomodulators were used to study immune pathways. We assessed brain c-Fos, brain-derived neurotrophic factor, and dendritic and glial cells by immunofluorescence, expression of neuroimmune genes by NanoString and performed brain proteomics.

**Results:** Bacterial monocolonization, conventionalization or administration of microbial products to germ-free mice altered mouse behavior similarly, acting through Toll-like receptor or nucleotide-binding oligomerization domain signaling. The process required CD11b^+^CD11c^+^CD103^+^ cell activation and migration into the brain. The change in behavior did not require the continued presence of bacteria and was associated with activation of multiple neuro-immune networks in the gut and the brain.

**Conclusions:** Changes in neural plasticity occur rapidly upon initial gut microbial colonization and involve innate immune signaling to the brain, mediated by CD11b^+^CD11c^+^CD103^+^ cell migration. The results identify a new target with therapeutic potential for DGBIs developing in context of increased gut and blood-brain barrier permeability.

**Highlights:**

- Microbiota impairment is a key factor in disorders of gut-brain interaction (DGBI)
- Microbial colonization induces changes in brain and behavior via innate immunity
- Microbial colonization activates multiple neuro-immune networks in gut and brain
- Behavioral change is mediated by CD11b^+^CD11c^+^CD103^+^ cells migration to the brain

## Introduction

Disorders of Gut-Brain Interaction (DGBI) encompass dozens of clinical conditions, including Irritable Bowel Syndrome (IBS), occurring in over 40% of adults and children that are related to any combination of gastrointestinal dysmotility, abdominal pain, immune dysregulation, gut dysbiosis and altered brain function^1,2^. Psychiatric comorbidities, mainly anxiety and depression, are common in patients with DGBI^3,4^. Although the gut microbiota has been implicated as a key player in the gut-brain axis^5^, the precise pathways driving abnormal brain function in these patients remain unclear. Thus, there is a gap in mechanistic pathway knowledge that needs to be addressed to better understand these complex disorders and to design treatments targeting their cause.

Gut microbiota profiles differ between patients with IBS and healthy controls^6^, which may stem from previous infectious gastroenteritis^7^, or combinations of life-long environmental factors that impact the microbiota, including long-term diet^8^, antibiotics^9^, or impaired initial seeding of maternal microbiota. Indeed, alteration of the maternal microbiota has been shown to alter offspring brain maturation, leading to abnormal behavior in adulthood^10–12^. Similarly, perturbation of the microbiota after birth, either due to stress or antibiotics, predisposes to dysfunction of the brain and gut. In a maternal separation model of IBS^13^, separating pups from their moms leads to intestinal dysbiosis, which is a critical determinant of the anxiety and depressive-like behaviors in adulthood, as these are not seen in germ-free mice^14^. Likewise, administration of antibiotics to pups results in long-term alterations in brain development and anxiety-like behavior, which can be prevented with probiotic supplementation^15^. In humans, a large population-based study showed that antibiotic treatment during infancy increases risk of development of mental illness in children^16^, all together suggesting that perturbation of the host-microbial crosstalk at a critical time of microbial colonization may have long-term consequences.

Despite the mounting evidence that microbiota shapes the neuroimmune system, detailed mechanisms underlying the initial phase of microbial-host interaction, are not fully understood. These studies are difficult to perform in humans, due to the complexity of microbiota-brain communication and high inter-individual variability in microbial profiles. Gnotobiotic technology offers a reductionist but controlled approach, allowing detailed studies of bacteria-host interactions at multiple timepoints, and concurrent investigation of behavior with neuroimmune pathways in the gut and the brain.

It is well established that compared with conventional mice, germ-free (GF) mice display anxiolytic and altered social behavior^17–19^, impaired blood brain barrier^20^, altered hypothalamic-pituitary-adrenal axis^21^ and impaired microglia function^22^, which normalize after bacterial colonization^19,21^. We took advantage of this knowledge and applied state of the art microbiology tools, combined with pharmacological and immune investigations to advance our mechanistic understanding of behavior and brain chemistry occurring after initial microbial colonization. Our studies identify a key role for microbially induced, CD11b^+^CD11c^+^CD103^+^ cell activation and migration that impacts brain neuroplasticity. The results can open new avenues for the development of therapies for DGBIs, particularly in context of infectious gastroenteritis or dysfunctional gut microbial colonization.

## Methods

### Gnotobiotic mice

Germ-free Swiss-Webster, C57BL/6, SCID and MyD88^−/−^ Ticam1^−/−^ mice were raised and maintained axenic at the Farncombe Family Axenic Gnotobiotic Unit (AGU), McMaster University, Canada. Conventionally raised Swiss-Webster mice were obtained from Taconic Farms (Hudson, NY, USA). Both sexes were included in this study. Handling of GF mice was carried out under the sterile conditional as previously described^23^. GF and colonization status was assessed regularly by direct bacteriology, immunofluorescence, and 16S gene PCR testing for culturable and unculturable organisms (for details see *Supplementary Methods*). All experiments were approved by the McMaster University Animal Care Committee.

### Bacterial colonization of germ-free mice

Germ-free mice (10-14 weeks old) were monocolonized with 10^9^ CFU of *E. coli JM83*, auxotrophic *E. coli HA107* or auxotrophic *Salmonella enterica* serovar *typhimurium* HA630 (STm^Aux^) via intragastric gavage. *E. coli HA107*, derived from *E. coli JM83*, only transiently colonizes (12-48 hours) mouse intestine^24^. *E. coli JM83* was gavaged once, and then the mice were gavaged 3 times weekly for 2 weeks with saline. The transient colonizers *E. coli HA107* and STm^Aux^ were gavaged 3 times weekly for 2 weeks. Additional GF mice were gavaged once with Altered Schaedler Flora (ASF,) or Specific Pathogen Free (SPF) microbiota and then 3 times weekly for 2 weeks with saline.

### Behavioral assessment

Mouse behavior was assessed in a custom designed isolator under gnotobiotic conditions using fully computerized system (Med Associates Inc, St. Albans, Vermont). Anxiety-like behavior was evaluated using light/dark preference test and depression-like behavior by the tail suspension test, as previously described^23^. At 2-, 4-, and 6-week time points, several mice (n=2-8) were sacrificed to verify the bacterial status as well as to obtain tissues.

### Pharmacological interventions

*E. coli 0111:B4* lipopolysaccharide (LPS, 2 mg/kg, Sigma, Canada) and Polyinosinic:polycytidylic acid (Poly I:C, 180 µg/ml, Sigma, Canada) were gavaged three times weekly for two weeks^24^. TLR4 inhibitor resatorvid, (TAK-242, MedChemExpress, United States) dissolved in 10% DMSO, 40% PEG300, 5% Tween-80 and saline, was administered daily via i.p at a dose of 5 mg/kg for two weeks. Cosalane (Southern Research Institute, Birmingham, AL) blocking Chemokine Receptor 7 (CCR7) activation and fingolimod (FTY720, GILENYA, Novartis, Canada) blocking immune cell migration, were administered three times weekly for two weeks. Cosalane, dissolved in 10% ethanol, was administered via i.p. injections in saline (2 mg/kg for 4 days and 1 mg/kg afterwards^25,26^). Fingolimod, dissolved in dimethyl sulfoxide, was added to the drinking water (0.4 mg/kg for 4 days and 0.2 mg/kg afterwards ^27^).

### Tissue processing

Brains were snap frozen in 2-Methylbutane over dry ice and stored at −80°C. Tissues were cut into 5μm sections and mounted onto frost Apex coated slides (Surgipath, Ontario, Canada) and stored at −20°C until processing. Region of interest in the brain included hippocampus (bregma −1.22mm and −2.70mm) and amygdala (bregma −1.22 and −2.18). Gut samples embedded in OCT were snap frozen in liquid nitrogen for storage at −80°C. Tissues were cut into 7μm cross sections and mounted onto Superfrost Plus coated slides (Fisher Scientific, Pittsburg, USA) and kept at −20°C until processing.

### Immunofluorescence staining

Brain sections were fixed in methanol, and proteins blocked using 10% normal serum with 1% BSA in TBS. Primary antibodies to BDNF and c-fos were applied overnight, and then visualized with fluorescent secondary antibodies. For dendritic cell, Iba1 and bacterial staining, brain sections were fixed in cold acetone and blocked with PBS containing 5% BSA. Brain sections were incubated overnight with anti-CD11b Alexa Fluor 488, anti-CD103 Alexa Fluor 594, anti-CD11c Alexa Fluor 647, anti-*E.coli*, anti-*S. typhimurium* or anti-Iba1 antibodies followed by application of appropriate secondary antibodies. For dendritic cell and bacterial staining, intestinal sections were fixed in 4% PFA, then incubated in blocking solution using PBS containing 5% FBS followed by PBS containing 10% rat serum. The slides were incubated overnight with anti-CD11b Alexa Fluor 488, anti-CD103 Alexa Fluor 555, anti-CD11c Alexa Fluor 647, anti-*E. coli* or anti-*S. typhimurium* antibodies followed by application of appropriate secondary antibodies.

Additional methodological details and list of antibodies are reported in *Supplementary Methods*.

### Gene expression study

RNA was extracted using the Rneasy Mini Kit and Dnase digestion during purification was carried out using the Rnase-free Dnase (both Qiagen Toronto, Canada). NanoString nCounter® Gene Expression CodeSet for mouse inflammation and a custom Nanostring gene codeset were run on colon and brain tissues, respectively, following manufacturer’s instructions (NanoString Technologies, Inc., Seattle, WA). The results were processed with nSolver 2.5 (NanoString Technologies), the Log2 ratios were then uploaded into Ingenuity Pathway analysis software (Qiagen) for further analysis. The network score is based on the hypergeometric distribution and calculated with the right-tailed Fisher’s exact test.

### Label-Free Quantitative Proteomics Assay using Nano-UPLC MS

Brain tissues were homogenized by sonication in 150μL of lysis buffer with 50mM ammonium bicarbonate, pH 8, 0.25% Rapigest SF (Waters), complete protease inhibitor (Roche), and phosphatase inhibitor cocktails I and III (Sigma-Aldrich). After centrifuge, supernatant protein concentrations were assayed by Bradford assay. To determine proteins expression, samples were diluted to 5 mg/mL with lysis buffer and reduced by incubation with 10 mM DTT. Then iodoacetamide was added to 20 mM, and samples were incubated at room temperature. Sequencing-grade modified trypsin (Promega) of 1:100 (w/w) was added to digest proteins samples for overnight. Next, samples were acidified to a final concentration of 1% TFA / 2% ACN and incubated at 60 °C for 2 hours. Finally, 50 μg of digested proteins per sample were transferred to Total Recovery LC Vials (Waters), and 50 fmol of MassPrep ADH standard (Waters) per μg of brain protein digests were added as an internal standard. Purified digested peptides were analyzed using reverse phase uplc (nanoacquity) with qtof ms/ms (Waters) as previously described^28^.

### Statistical analysis

Statistical analysis was performed using Prism 6 software. Statistical comparisons were performed using one-way ANOVA followed by Dunn’s test for multiple comparison, student t-test, Mann-Whitney t-test, or Wilcoxon’s signed rank t-test as appropriate. Benjamini and Hochberg FDR correction method was used when multiple comparisons were performed for NanoString. A p-value of <0.05 was considered statistically significant.

## Results

Germ-free Swiss-Webster mice were assessed before and after colonization with Specific Pathogen Free (SPF) microbiota or the Altered Schaedler Flora (ASF) (**Fig. 1A**), conventionally raised mice (permanent conventional, pcSPF) were used as controls. Compared to pcSPF mice, germ-free mice displayed more exploratory (anxiolytic) behavior spending more time in the illuminated compartment during the light preference test and exhibited less depressive behavior being more mobile during the tail suspension test (**Fig. 1B**). Ex-germ-free mice colonized with ASF or SPF microbiota displayed similar behavior as pcSPF mice in the light-preference and tail-suspension tests (**Fig. 1B**) at 2 and 4 weeks post-colonization. Overall locomotor activity was similar between groups, ruling out sickness behavior in colonized mice (**Supplementary Fig. 1A**). Control experiments, where germ-free mice were repeatedly gavaged with sterile saline, verified that handling itself did not affect mouse behavior (**Fig. 1C**). Next, we tested whether a single bacterial strain alters mouse behavior using non-pathogenic strain *Escherichia coli JM83,* known to induce immune system maturation^24^. Germ-free Swiss-Webster mice mono-colonized with *E. coli JM83* spent less time in the light compartment and displayed more immobility at 2- and 4-weeks post-colonization compared to before colonization (**Fig. 1D**).

**Figure 1:**
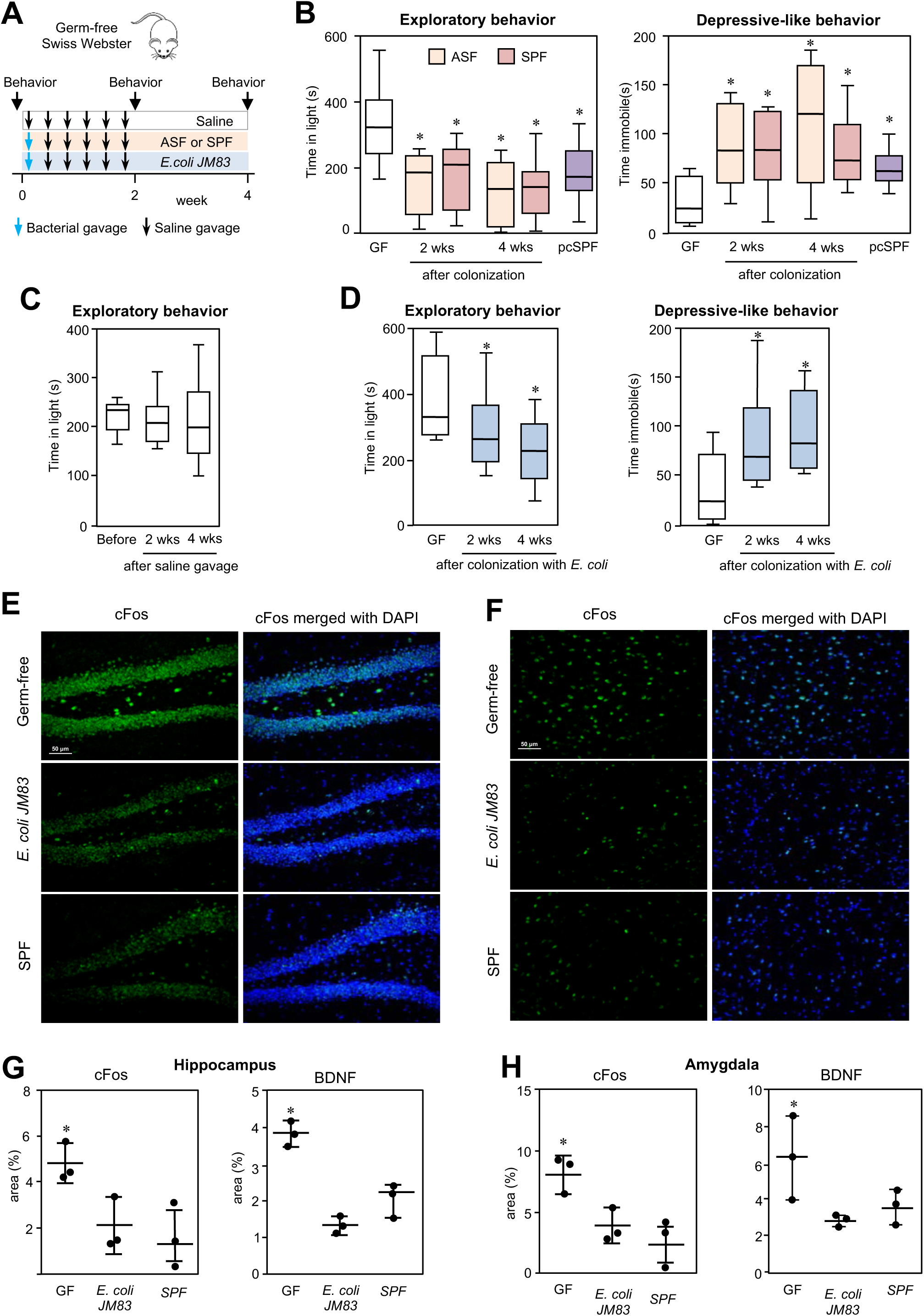
Colonization with E. coli JM83 alters behavior similarly to colonization with ASF and SPF microbiota. A) Experimental design. B) Exploratory behavior (using the light preference test) and depressive-like behavior (using the tail suspension test) of germ-free (GF) mice before and after colonization with ASF and SPF microbiota, compared to conventionally raised mice (pcSPF) (n=16/group). C) Exploratory behavior of GF mice before and after saline gavage (n=8). D) Behavior of GF mice before and after colonization with *E. coli* JM83 (n=18). E) Hippocampus c-fos expression in GF, *E. coli* mono-colonized and SPF mice. F) Amygdala c-fos expression in GF, *E. coli* mono-colonized and SPF mice. G) Quantification of c-fos and BDNF positive areas in hippocampus. H) Quantification of c-fos and BDNF positive areas in amygdala. Statistics by Wilcoxon’s signed rank test and ANOVA, *p<0.05 vs GF.

There was no evidence of sickness behavior (**Supplementary Fig. 1B**) or measurable inflammation compared to GF controls (*data not shown; below detection*). Using immunofluorescence, we assessed c-fos and brain-derived neurotrophic factor (BDNF) expression as markers of neural activation and plasticity, respectively. Level of c-fos and BDNF were lower in the hippocampus and amygdala (**Fig. 1E-H**, **Supplementary Fig. 1C,D**), two regions involved in control of exploratory/anxiety and depressive-like behaviors, in mice colonized with SPF microbiota or *E. coli* compared to germ-free mice. Thus, bacterial colonization with both diverse and highly restricted microbiota, or even a single bacterial strain, normalizes behavior and brain chemistry of germ-free mice.

To determine whether normalized behavior requires the continuous presence of bacteria, germ-free Swiss-Webster mice were gavaged for 2 weeks with a transient colonizer *E. coli HA107,* a triple auxotrophic mutant derived from *E. coli JM83*^24^ (**Fig. 2**). The reversion to a germ-free status was confirmed by negative cultures of mouse stool and cecum contents at 4 and 6-weeks post-colonization. At 2 weeks, *E. coli HA107*-colonized mice displayed lower exploratory behavior that remained unchanged at weeks 4 and 6, with mice behaving similarly as mice colonized with *E. coli JM83* (**Fig. 2**).

**Figure 2:**
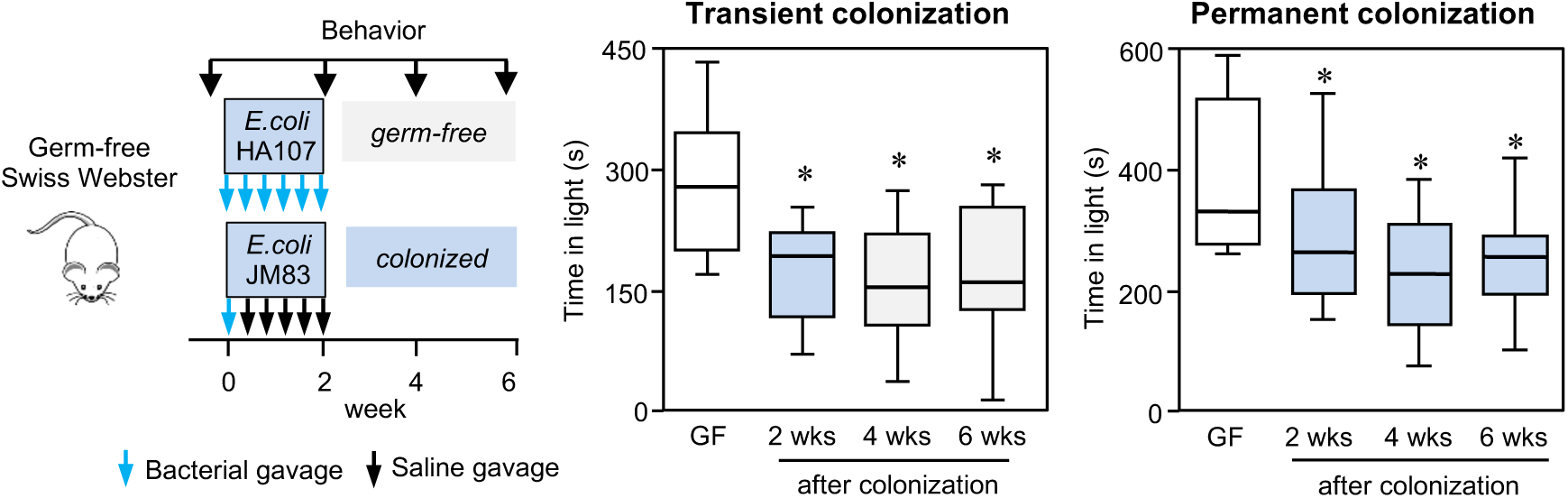
Transient bacterial colonization induces long-term changes in exploratory behavior. Behavior of germ-free (GF) mice (n=20) before and after colonization with permanent colonizer *E. coli* JM83 and transient colonizer *E. coli* HA107. Statistics by Wilcoxon’s signed rank test, *p<0.05 vs GF.

To investigate the putative mechanisms underlying this pronounced and persistent change in behavior, we assessed behavior of germ-free C57BL/6, MyD88^−/−^ Ticam1^−/−^ and SCID mice before and after colonization with *E. coli JM83* (**Fig. 3A**). Knowing that change in behavior is already present after 2 weeks, we decided to investigate mouse behavior at 1 and 3 weeks post-colonization. Similar to Swiss Webster mice, mono-colonized C57BL/6 mice displayed less exploratory behavior and increased immobility. While SCID mice behaved similarly as C57CL/6 mice, MyD88^−/−^ Ticam1^−/−^ mice did not change their behavior after colonization (**Fig. 3A**). In parallel, brain BDNF and c-fos expression were similar in germ-free and mono-colonized MyD88^−/−^Ticam^−/−^ mice (**Supplementary Fig. 2A,B**), all together suggesting that innate, but not adaptive immunity mediates the post-colonization behavior changes.

**Figure 3:**
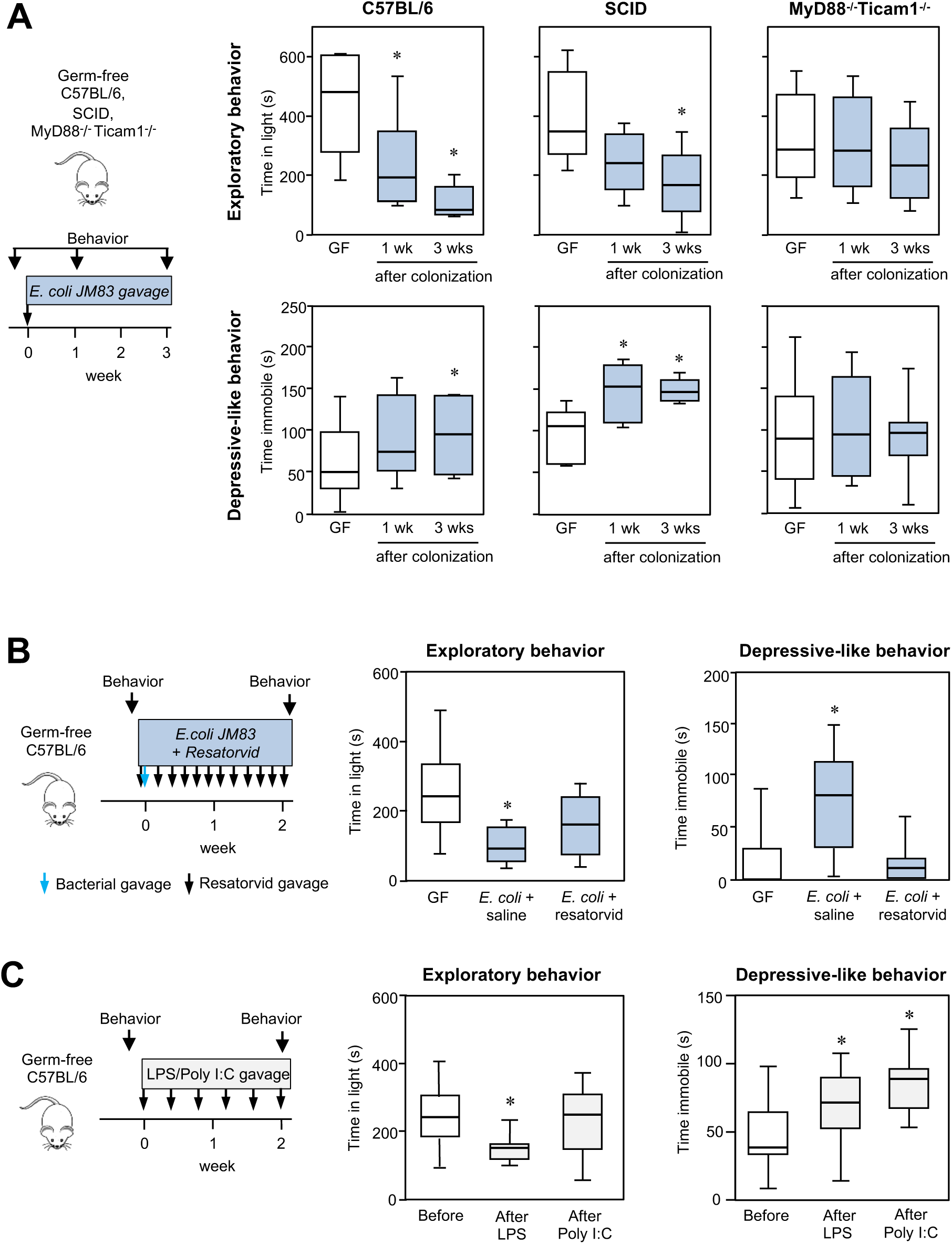
MyD88/Ticam and TLR4 signaling is crucial for the behavioral changes after microbial colonization. A) Behavior of germ-free (GF, n=14), SCID (n=10), and MyD88^−/−^Ticam1^−/−^ (n=18) mice before and after colonization with *E. coli* JM83. B) Behavior of GF mice (n=7) before and after colonization with *E. coli* JM83 and treatment with TLR4 inhibitor (Resatorvid). C) Behavior of GF mice (n=8) before and after repeated LPS or poly I:C gavage. Statistics by Wilcoxon’s signed rank test, *p<0.05 vs GF.

To confirm the role of TLR signaling, we gavaged germ-free C57BL/6 mice with TLR-4 inhibitor resatorvid before and after colonization with *E. coli JM83.* Compared to saline, resatorvid administration prevented changes in exploratory and depressive behaviors (**Fig. 3B**). To determine whether normalization of behavior can be induced solely by activation of TLRs, in the absence of microbiota, germ-free mice were gavaged repeatedly with *E. coli* LPS, a TLR4 agonist, or poly I:C, a TLR3 agonist (**Fig. 3C**). LPS induced similar behaviors as mono-colonization, while poly I:C changed behavior only in the tail-suspension test. This likely reflects differences in the magnitude of immune stimulation as both TLR4 and TLR3 expression increased in the jejunum of LPS and poly I:C gavaged mice (10- and 2-fold, respectively, **Supplementary Fig. 2C,D**) while in the colon, only TLR4 expression increased (2-fold).

When investigating which innate immune cells mediate this process, we focused on intestinal dendritic cells (DCs), as main antigen processing cells at the microbial-host interface. We assessed behavior before and after 2 weeks of mono-colonization with *E. coli JM83* in germ-free mice treated with cosalane, inhibitor of CC-Chemokine Receptor 7 (CCR7)^25,26^, or with sphingosine 1-phosphate receptor modulator fingolimod^27^. While fingolimod blocks migration of several types of innate immune cell, cosalane inhibits DCs migration in response to CCL19 and CCL21 agonists. Whilst saline-treated mice normalized their behavior after colonization, we found no change in exploratory or depressive-like behavior in cosalane- or fingolimod-treated mice (**Fig. 4A**); neither of these mice displayed sickness-like behavior (**Supplementary Fig. 2E**).

**Figure 4:**
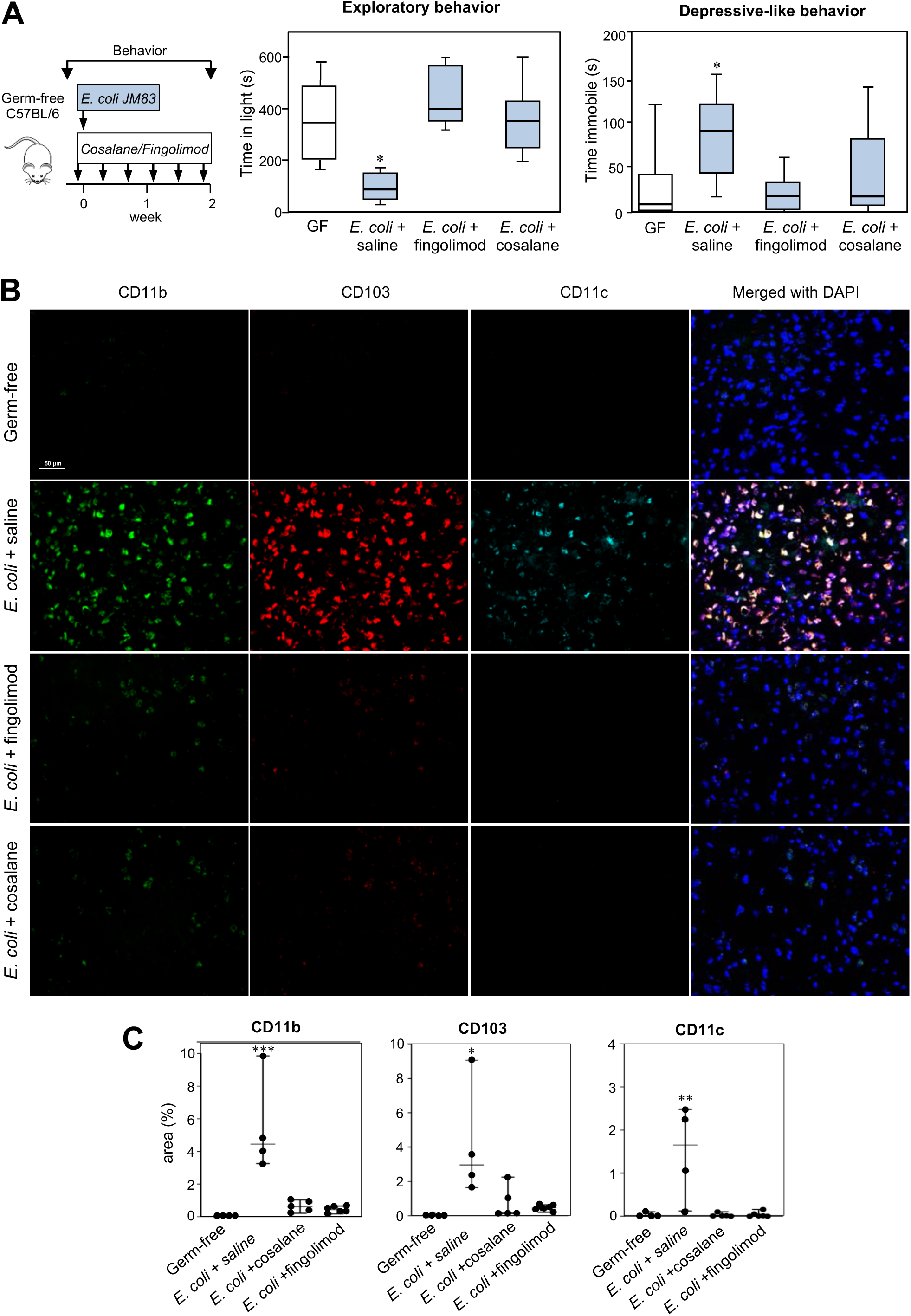
Blocking dendritic cells prevents changes in behavior after bacterial colonization. A) Behavior of germ-free (GF) C57BL/6 mice before and after 2 weeks of colonization with *E. coli* JM83 treated with saline, cosalane or fingolimod (Wilcoxon’s signed rank test; n=7/group). B) CD11b, CD103 and CD11c expression in the amygdala of GF and mono-colonized mice treated with saline, cosalane or fingolimod. C) Quantification of CD11b, CD103 and CD11c positive area in amygdala of GF and mono-colonized mice treated with saline, cosalane or fingolimod. Statistics by ANOVA, *p<0.05, **p<0.01, ***p<0.001 vs GF mice.

We then assessed expression of CD11b, CD11c and CD103 as markers of intestinal DCs by immunofluorescence. Although there was no difference in CD11b and CD11c staining in the intestine, CD103 expression was higher in mono-colonized mice treated with saline compared with germ-free, or cosalane- and fingolimod treated mice (**Supplementary Fig. 3A,B**). In the brain, no CD11b^+^CD103^+^CD11c^+^ cells were present in germ-free mice, but they were found in high numbers in *E. coli*-monocolonized mice, both in the amygdala (**Fig. 4B,C**) and hippocampus (**Supplementary Fig. 3C,D**). Importantly, treatment with both fingolimod and cosalane reduced brain CD11b^+^CD103^+^CD11c^+^ cells almost to levels observed in germ-free mice. To further characterize these cells, we performed Iba1 immunostaining, specific for macrophages and microglial cells, in the brain and found no overlap between CD11b and Iba1 positivity (**Supplementary Fig. 4**), suggesting these cells were DCs.

To link CD11b^+^CD11c^+^CD103^+^ cells in the brain with the intestinal bacteria, we used immunofluorescence staining specific for *E. coli* and found occasional presence of bacterial fragments within CD11b^+^CD11c^+^CD103^+^ DCs in the small intestine of monocolonized mice (**Fig. 5A**), and in rare occasions in CD11b^+^CD11c^+^CD103^+^ cells the hippocampus or amygdala (**Fig. 5B**), suggesting that DCs can migrate from the intestine and transport bacterial fragments into the brain.

**Figure 5:**
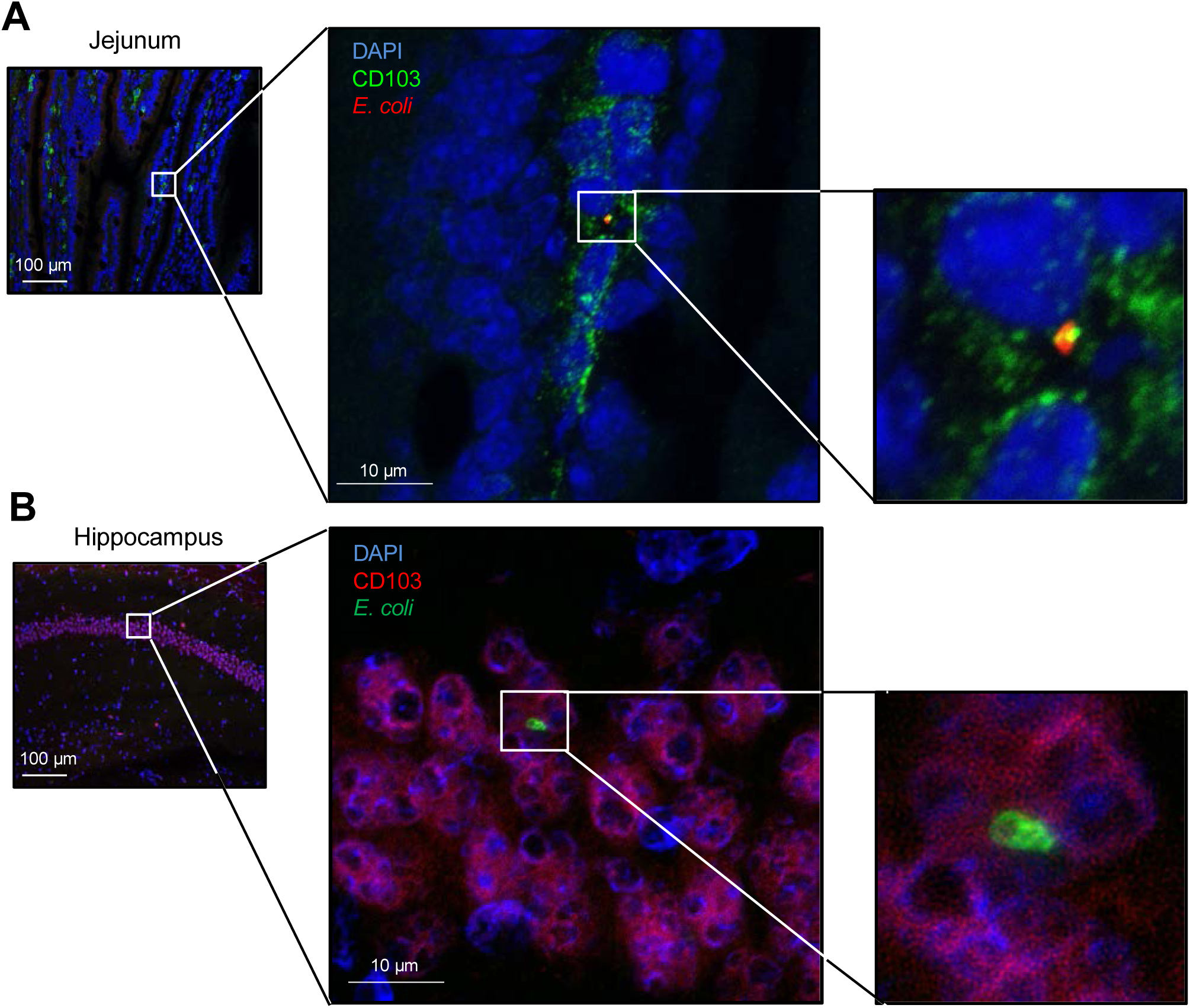
Presence of E. coli immunoreactivity within CD103^+^ cells in the intestine and brain. A) Representative images of immunofluorescent staining of CD103^+^ cells (green) and *E. coli*^+^ immunoreactivity (red) in the jejunum of germ-free (GF) mice mono-colonized with *E. coli* JM83. White squares indicate the magnified areas. B) Representative images of immunofluorescent staining of CD103^+^ cells (red) and *E. coli*^+^ immunoreactivity (green) in the hippocampus of GF mice mono-colonized with E. coli JM83. White squares indicate the magnified areas.

To confirm these findings, and further explore the role of the innate immune system, we colonized germ-free MyD88^−/−^ Ticam1^−/−^ mice with the transiently colonizing auxotrophic strain *Salmonella enterica* serovar *Typhimurium* (STm^Aux^), previously shown to induce robust adaptive immunity in these mice through NOD1/NOD2 nodosome signaling^29^. Colonization with STm^Aux^ decreased exploratory behavior in MyD88^−/−^ Ticam1^−/−^mice (**Supplementary Fig. 5A**) to a similar degree as previously seen in C57BL/6 mice colonized with *E. coli*, but it did not affect their overall locomotor activity. Using immunofluorescence, we found STm^Aux^ bacterial fragments within CD11b^+^CD11c^+^CD103^+^ DCs in the intestine (**Supplementary Fig. 5B**), and occasionally bacterial fragments within CD11b^+^CD11c^+^CD103^+^ cells in the brain (**Supplementary Fig. 5C**). All together, these data suggest that the intestinal DCs, activated either through TLR or NOD signaling, are crucial for the establishment of normal mouse behavior after initial bacterial colonization.

To further investigate neuroimmune mechanisms associated with the behavioral normalization, we assessed colonic gene expression in Swiss Webster, C57BL/6, and MyD88^−/−^ Ticam1^−/−^ mice, before and after colonization with *E. coli*. Mono-colonization altered expression of most of the 250 immune genes studied (**Supplementary Fig. 6A**). To identify the relevant pathways, we searched for genes that fulfilled the following criteria: 1) change in transcripts at 2 weeks post-colonization compared to GF status and 2) similar expression at 2- and 4-weeks in C57BL/6 mice; 3) unchanged expression in MyD88^−/−^ Ticam^−/−^ mice after colonization. These criteria were chosen as behavior of immunocompetent mice changed dramatically at 2 weeks post-colonization with no further alteration at 4 weeks, while no behavior changes were observed in MyD88^−/−^ Ticam^−/−^ mice. The criteria were fulfilled by 26 genes, mainly related to the innate immunity (**Fig. 6A**). Ingenuity pathway analysis revealed that many were related both to immune function and neural development and behavior (**Fig. 6C**). The main canonical pathways were the acute phase response (P=1.71*10^−32^), toll-like receptor (P=8.11*10^−26^), pattern recognition receptors (P=4.38*10^−25^) and glucocorticoid receptor (P=1.26*10^−19^. Several genes were previously reported to be involved in _the formation and maintenance of_ neuronal plasticity, including the Rho GTPases *RhoA* and *Cdc42*^30^, *CCR2*^31^, *MEF2* and members of p38 MAPK pathway^32,33^, or linked to glucocorticoid receptors, whose priming determines formation of learning-dependent synapses^34^.

**Figure 6:**
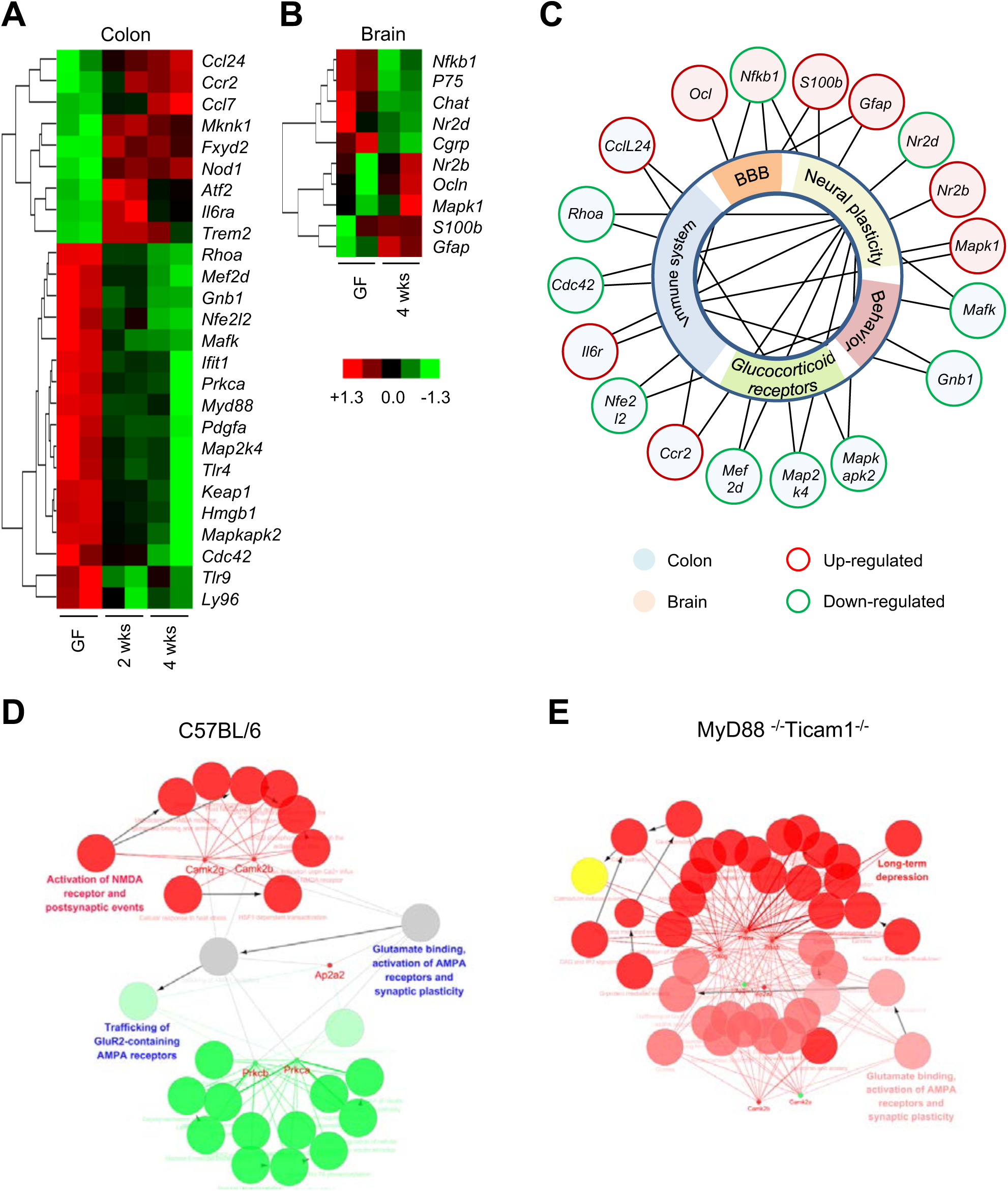
Multiple neuro-immune genes and protein networks are altered by bacterial colonization. A) Colonic immune genes involved in altered behavior in mono-colonized Swiss Webster mice. B) Brain neuro-immuno-epithelial genes altered by mono-colonization in Swiss Webster mice. C) IPA network of brain and colonic genes altered by bacterial mono-colonization in Swiss Webster mice. D,E) Schematic illustration of systematic rewiring of dynamic biological networks related to neural plasticity induced by bacterial mono-colonization in C57BL/6 mice (D) and MyD88^−/−^ Ticam1^−/−^ mice (E). The expression of each protein is visualized with small dot (green: increase, red: decrease). Biological pathways and functions are visualized as circular nodes linked with related proteins based on their change of expression: up-regulated: green, down-regulated: red.

To link the intestinal immune profile with the behavioral change, we assessed brain gene expression using a custom NanoString codeset (72 neuro-immuno-epithelial genes, **Supplementary Fig. 6B**). Colonization affected 10 genes associated with neuronal development and blood-brain barrier function (**Fig. 6B,C**), with the canonical pathways of the toll-like receptor (P=4.34*10^−9^) and axonal guidance signaling (P=5.25*10^−4^). Altered expression of *Ocln*, *GFAP* and *ERK2* imply remodeling of blood-brain barrier^20^, neuroglial activation^35^ and altered neuronal plasticity^33^, respectively. We also found phenotypic switch in glutamate NMDA receptors with down-regulation of *NR2D* and up-regulation of *NR2B*, the 2 subunits with different functional properties^36^. *NR2D*, with longer deactivation time and lower NMDA-mediated currents, is abundant in the embryonic brain of conventionally raised mice but its levels abruptly decrease first week after birth ^37^, possibly linked to initial bacterial colonization post-partum.

We then performed brain proteomic analysis using Nano-UPLC MS. Colonization altered 97, 163 and 149 proteins in Swiss Webster, C57BL/6 and MyD88^−/−^ Ticam1^−/−^ mice, respectively, predominantly G-proteins (**Supplementary Table 1**). The main neural pathways included glutamate binding, activation of AMPA receptors and synaptic plasticity (**Fig. 6D,E**). While the changes in protein expression were similar in Swiss Webster and C57BL/6 mice, they differed in MyD88^−/−^ Ticam1^−/−^ mice, suggesting key role of TLR signaling in the development of neural plasticity (**Fig. 6D,E and Supplementary Fig. 9**). From the altered proteins, ten were directly linked to neural plasticity: Protein kinase C^38^ (Prkca, Prkcb, Prkcg), Calcium/calmodulin-dependent protein kinase (Camk2a) and Adaptor protein complex 2 (Ap2a1, Ap2a2) are involved in glutamate binding, activation of AMPA receptors and synaptic plasticity^39^ (R-MMU:399721, P<0.05), while Camk2b and Ras-related protein (Rab8a) are implicated in long-term synaptic plasticity^40^ (GO:0048169, P<0.05). These neural system changes were associated with altered levels of proteins linked to the immune system, with the pathways most affected being antigen processing, as well as myeloid cell differentiation, regulation of immune response, and T cell activation.

## Discussion

Under physiological conditions, gut microbial colonization occurs postnatally^41^ and is associated with changes in the immune and neural systems^5,42,43^, many of which are reproduced in germ-free mice colonized with microbiota, both at early life^42,43^ or adulthood^17–21^. However, the exact pathways underlying microbiota-gut-brain axis communication at first encounter with microbiota remain largely unknown. In this study we found that *de novo* microbial gut colonization activated innate immune system pathways through TLR or NOD signaling. This was associated with activation and migration of intestinal DCs, initiating neuro-immune signaling within the gut-brain axis that culminated in changes in brain plasticity and mouse behavior maturation. The immune mediated changes in behavior were induced by both complex communities or single bacterial strains, and were also achieved in the absence of bacteria, by stimulating pattern recognition receptors. The changes in behavior induced post-colonization were long-lasting and did not require the continuous presence of gut bacteria, as demonstrated by experiments using transient bacterial colonizers in which mice reverted to their original germ-free status.

Although adaptive immune responses have been shown to affect brain function in conventional adult mice^44^, we demonstrate here that innate immunity, either through MyD88-dependent pathways downstream from TLRs or through NOD signaling, play a key role in shaping behavior following bacterial colonization. This is in agreement with studies showing that bacterial products through TLR4 signaling impact social behavior and dopaminergic pathways^45,46^ and that NOD signaling modulates stress-induced anxiety coupled with changes in the serotonergic system^47^.

In the gut, DCs are prime regulators of the innate immune system, shaping tolerance or immune responses to microbial and food antigens^48^. *Lamina propria* DCs are activated by bacteria via TLR-MyD88 or NOD signaling, migrating to mesenteric lymph nodes, with capacity to cross the blood-brain-barrier under certain conditions^49,50^. We show that pharmacological block of DC prevented behavioral adaptations to colonization, accompanied by absence of CD11b^+^CD103^+^CD11c^+^ cells in the brain areas involved in exploratory and depressive-like behavior. The CD11b^+^CD103^+^CD11c^+^ cells in brains of colonized mice expressed intracellular immunoreactivity for bacteria, which in a combination of markers typical for intestinal DCs, strongly support their gut origin. Overall, the results suggest a crucial role for innate immunity and intestinal DCs in shaping normal function of the neural system and brain upon initial gut colonization.

We found that colonization affected multiple neuro-immune pathways in the gut and the brain, including the canonical pathway of glucocorticoid receptor which regulates emotion and cognition^51^. Several genes directly involved in neuronal plasticity were altered by bacterial colonization, including phenotypic switch in NMDA receptors subunits, which in conventional mice appears postnatally^37^. Proteomic analysis confirmed that colonization had a major impact on several brain neuro-immune networks involved in glutamate binding and activation of AMPA receptors that play a key role in neuronal plasticity^52,53^, as well as ten specific proteins directly involved in the induction and the maintenance of synaptic plasticity^38–40^. These data show that behavior adaptations after gut colonization associate with major changes in brain chemistry and function, which are immune driven. Moreover, the altered behavioral phenotype persists even after mice revert to germ-free status, as shown in experiments using transient bacterial colonizers.

It is well established that behavior and brain chemistry differ between germ-free and conventional mice^17,19,54^. In agreement with previous studies that showed that germ-free mice display anxiolytic behavior compared with conventional mice^17,19^, we found that bacterial colonization had both anxiety- and depressive-like effect as evidenced by mice spending consistently less time in the illuminated compartment, and displaying an increased despair behavior as assessed by the tail suspension test. Our data suggest that bacterial colonization alters multiple facets of behavior, in part leading to different risk assessment and a more “cautious” behavior phenotype, which may provide evolutionary benefit for mouse survival. Thus, what we consider “normal behavior” in healthy conventional mice, whether in terms of exploration, coping with novel situation and risk-assessment^55^, is determined by initial microbial-immune interactions.

Impairment of this process due to abnormal maternal microbiota seeding^10–12^ or perturbation of the microbiota after birth^13,14^ with aberrant immune response, could predispose to anxiety, depression or generalized psychological vulnerability^56^, and lead to the gut-brain dysfunction and mental health disorders^23^. During adulthood, severe gastroenteritis associated with increased epithelial, endothelial and blood-brain barrier permeability could activate these pathways^57,58^ and worsen pre-existing gut-brain dysfunction, which is a known risk factor for post-infective IBS^7,59^, or contribute to development of new DGBIs^7^.

Our study has several limitations. It is difficult to assess changes in neural functioning, especially behavior, at the early postnatal period and therefore we studied the effects of initial bacterial colonization in young adult germ-free mice. To dissect the underlying mechanisms, we used a reductionist approach involving bacterial monocolonization of immunocompetent and immunodeficient mice. As controls, we used conventionally raised mice (permanent conventional, pcSPF), demonstrating that monocolonization resulted in similar behavior as observed in pcSPF mice. This approach allowed us to study behavior in individual mice before and at different time points post colonization, instead of comparing separate cohorts of mice. We identified DCs with typical intestinal markers in the brain, some containing bacterial fragments, strongly supporting their intestinal origin. However, we did not use a dedicated technique, such as a light convertible mouse reporter line^60^, to track direct migration of intestinal DCs into the brain. This would be extremely difficult to implement in our germ-free set-up, where all experiments are performed within custom made gnotobiotic behavioral isolators.

In conclusion, our study identifies activation of the intestinal innate immune system as the first step initiating maturation of behavior upon bacterial colonization, and strongly suggest a key role of DCs as specific cells involved in microbiota-gut-brain communication. Our findings may have bearing on disorders of gut-brain interactions and psychiatric diseases, in which altered innate immune signaling has been implicated^61,62^.

## Funding

Canadian Institutes of Health Research (CIHR) Foundation grant #143253 (PB, SMC) National Institute of Health (NIH) grant #R33 MH108167 (PB)

SH and OPS were supported by grants from the Swiss National Science Foundation (SNSF; grants 169791 and 200382).

## Abbreviations used in this paper

DGBI: Disorders of Gut-Brain Interaction
IBS: Irritable Bowel Syndrome
GF: Germ-free
ASF: Altered Schaedler Flora
SPF: Specific Pathogen Free
DC: Dendritic cell
TLR: Toll-Like Receptor
NOD: Nucleotide-binding Oligomerization Domain
CCR7: Chemokine Receptor 7
BDNF: Brain-Derived Neurotrophic Factor
LPS: Lipopolysaccharide
SCID: Severe Combined ImmunoDeficiency
PBS: Phosphate-Buffered Saline
TBS: Tris-Buffered Saline
BSA: Bovine Serum Albumin

## Conflicts of interest

The authors disclose no conflicts.

## Authors contributions

Conceptualization: PB, SMC, VP, NK, EFV, FC; Methodology: VP, NK, HZ, CS, PB, KDM, SH, MGS, OPS; Software: HZ, CS; Validation: VP, NK, JL, GDP; Formal analysis: VP, NK, HZ, CS, PB; Investigation: VP, NK, HZ, JL, GDP, PB; Resources: EFV, KDM, SH, AJM, MGS, FL, OPS; Writing – original draft: VP, NK, PB; Writing – review & editing: VP, NK, PB, GDP, CS, FC, EFV, KDM, SMC, SH; Visualization: VP, NK, PB; Funding acquisition: PB, SMC, SH, OPS; Project administration: VP, PB; Supervision: PB, SMC.

## Data and materials availability

NanoString data have been deposited in the Gene Expression Omnibus (GEO) database with accession code (GSE138321). The mass spectrometry proteomics data have been deposited to the ProteomeXchange Consortium via the PRIDE [1] partner repository with the dataset identifier PXD016152. The authors declare that all other relevant data supporting the findings of this study are available within the paper and its Supplementary Information files, or from the corresponding author on request.

## Acknowledgments

The authors are very grateful to Dr. M. Suto and Mrs. K. Sides from Southern Research Institute, Birmingham, AL, USA, for providing cosalane. PB holds the Richard Hunt-Astra Zeneca Research Chair in Gastroenterology. The authors thank Dr. Heather Galipeau and the McMaster’s AGU staff for their support with gnotobiotic experiments, and the staff at the Centre for Advanced Light Microscopy (CALM) for support in using the confocal microscopes.

## Supplementary Methods

### Gnotobiotic mice

Germ-free Swiss-Webster, C57BL/6, SCID and MyD88^−/−^ Ticam1^−/−^ mice were raised and maintained axenic in sterile isolators at the Farncombe Family Axenic Gnotobiotic Unit (AGU) of the Central Animal Facility, McMaster University, Canada. Conventionally Swiss-Webster mice (pcSPF) were obtained from Taconic Farms (Hudson, NY, USA). Both sexes were included in this study. To ensure sterility, handling of GF mice was carried out under the most scrupulously clean conditions as previously described ^23^. All mice were maintained on a 12-hour day/night cycle with unrestricted access to food and water. GF and mono-colonization status was assessed regularly by direct bacteriology, immunofluorescence and 16S gene PCR testing for culturable and unculturable organisms for ASF and SPF microbiota colonized mice. All experiments were approved by the McMaster University Animal Care Committee.

### Bacterial colonization of germ-free mice

Germ-free mice (10 - 14 weeks old) were mono-colonized with 10^9^ CFU of *E. coli JM83*, *E. coli HA107* or auxotrophic *Salmonella enterica* serovar Thyphimurium HA630 (STm^Aux^) via intragastric gavage. *E. coli HA107* strain is a mutant form of the parental strain *E. coli JM83*, which lacks the ability to synthesize *meso-diaminopimelic acid (m-DAP)* and *D-isomer of alanine (D-Ala)* required in the peptidoglycan crosslink of the cell wall and thus only transiently colonizes (12-48 hours) mouse intestine ^24^. The permanent colonizer *E. coli JM83* was gavaged once, and then the mice were gavaged 3 times weekly for 2 weeks with normal saline. The transient colonizers *E. coli HA107* and STm^Aux^ were gavaged 3 times weekly for 2 weeks. Mono-colonized mice were maintained in sterile isolators within the AGU. Additional groups of GF mice were gavaged once with restricted Altered Schaedler Flora (ASF, consisting of 8 bacterial strains) or diverse Specific Pathogen Free (SPF) microbiota and then gavaged 3 times weekly for 2 weeks with normal saline. These mice were housed in ultraclean conditions using ventilated racks. *E. coli JM83* was grown in LB broth and *E. coli HA107* was grown in D-Ala (200μg/ml)/m-DAP (50μg/ml)-supplemented LB broth and incubated with shaking at 160 rpm at 37°C for 12 hours. Bacteria were harvested by centrifugation (15 min, 3500X g) in a 400ml sterile flask, washed in sterile PBS and concentrated to a density of 10^9^ CFU/ml in PBS.

Auxotrophic *S. typhimurium* HA630 (STm^Aux^) which only transiently colonizes mouse intestine was grown in D-Ala (200μg/ml)/m-DAP (50μg/ml)-supplemented LB broth and incubated shaking at 150 r.p.m., at 37 °C for 16 h followed by a second incubation in 500 mL fresh medium under the same conditions for 15 more hours. Bacteria were harvested by centrifugation (15 min, 4816X *g*, 4 °C), washed twice with cold PBS, and resuspended to a density of 10^9^ CFU/ml, as previously described^29^. All the bacterial preparations were performed aseptically under a sterile laminar flow hood. The bacterial suspensions were sealed in sterile tubes, with the outside surface kept sterile, and imported into sterile isolators, where 200 μl (10^9^ CFU) were gavaged into germ-free mice.

### Behavioral assessment

Mouse behavior was assessed in a custom designed isolator under gnotobiotic conditions using fully computerized system (Med Associates Inc, St. Albans, Vermont). Anxiety-like behavior was evaluated using light/dark preference test as previously described ^23^. Briefly, each mouse was placed in the center of an illuminated box connected to a smaller dark box, and its behavior was recorded for 10 min. Total time spent in the illuminated area, total distance travelled and average velocity were assessed. Depression-like behavior was assessed using the tail suspension test as previously described ^23^. Briefly, each mouse was suspended by its tail for 5 min and the time it spends immobile was recorded. An animal was considered immobile when it did not show any body movement and hanged passively.

At the end of 2-, 4-, and 6-week time points, several mice (n=2-8) were sacrificed to verify the bacterial status as well as to obtain tissues (brain, blood, colon, small intestine) in order to run various assays.

### Gnotobiotic husbandry and microbial analysis

Isolators that house GF mice undergo meticulous protocols to prevent introduction of microbes from animal handlers or the environment. Samples of the imported materials are taken for aerobic and anaerobic bacterial culture regularly. Feces and bedding are taken from the isolator for direct bacteriology, microscopy and 16S gene PCR testing of intestinal contents to test for culturable and unculturable organisms. Behavioral experiments on GF mice colonized with single or multiple bacteria were carried out in smaller ‘surgical’ isolators separate from the breeding isolator to minimize risk of bacterial contamination of the stock during manipulations. Cecal contents from mono-colonized mice were suspended and serially diluted in sterile 1X PBS and plated on LB agar plate for 48 hours. Cecal contents from mice treated with *E. coli HA107* strain were incubated in LB plate supplemented with m-DAP and D-Ala. To confirm *de novo* GF status after colonizing with *E. coli HA107*, cecal contents were plated in supplemented LB agar plate 2 and 4 weeks after the last gavage. Bacterial immunofluorescence (cat no. L7005, ThermoFisher, Canada) staining of the cecal contents was carried out to ensure the sterility of the GF mice.

### Lipopolysaccharide (LPS), Polyinosinic:polycytidylic acid (Poly I:C) and TRL4 inhibitor

Sterile *E. coli 0111:B4* LPS (2 mg/kg, Sigma, Canada) and Poly I:C (180 µg/ml, Sigma, Canada) were gavaged into germ-free C57BL/6 mice three times weekly for two weeks ^24^. Mice treated with TLR4 inhibitor (Resatorvid, TAK-242, MedChemExpress, United States) were colonized once with the permanent colonizer *E.coli JM83*. Resatorvid was dissolved in 10% DMSO, 40% PEG300, 5% Tween-80 and 45% physiological saline to a final concentration of 1.25 mg/ml. Resatorvid was administered to mice once a day via i.p injections diluted in saline solution at a dose of 5 mg/kg for two weeks. Behavior was assessed before and after treatment.

### Cosalane and fingolimod drug treatment

Following the unique gavage with the permanent colonizer *E.coli JM83* the mice were treated by saline solution, by cosalane drug (obtained from the Southern Research Institute, Birmingham, AL) blocking CCR7 (Chemokine Receptor) activation or fingolimod drug (FTY720, GILENYA, Novartis, Quebec, Canada) blocking CCR7 activation and DCs migration. Drug administration was performed three times weekly for two weeks. Cosalane was stabilized in 10% ethanol and fingolimod was diluted in DMSO (dimethyl sulfoxide). Both drug solutions were prepared and filtered in sterile conditions. Cosalane was administered to mice via i.p. injections diluted in saline solution (2 mg/kg in the 4 first days and then 1 mg/kg for the next days to prevent any toxic effect on the mice ^25,26^). Fingolimod was added to the autoclaved drinking water (0.4 mg/kg in the 4 first days and then 0.2 mg/kg for the next days to prevent any toxic effect on the mice ^27^). Behavior was assessed before and after treatment.

### Brain tissue processing

Upon sacrifice of mice, brains were immediately snap frozen in 2-Methylbutane (isopentane) over dry ice and stored at −80°C. All tissues were then cut into 5μm sections on a Microm HM 550 cryostat (Thermo Scientific, WI, USA) and mounted onto pre-cleaned double frost Apex coated slides (Surgipath, Ontario, Canada) and stored at −20°C until processing. Region of interest in the brain included hippocampus (sections between bregma −1.22mm and −2.70mm) and amygdala (sections between bregma −1.22 and −2.18).

### Intestinal tissue processing

Upon sacrifice of mice, pieces of jejunum were embedded in optimal cutting temperature (OCT) compound and immediately snap frozen in liquid nitrogen for storage at −80°C. All tissues were then cut into 7μm cross sections on a Microm HM 550 cryostat (Thermo Scientific, WI, USA) and mounted onto pre-cleaned Superfrost Plus coated slides (Fisher Scientific, Pittsburg, USA) and kept at −20°C until processing.

### Immunofluorescence staining

Brain sections were allowed to thaw at room temperature for one hour then washed in phosphate-buffered saline (PBS, pH 7.4) for 5 min to remove any residual OCT. Tissue sections were fixed in methanol for 10 min, and proteins were blocked using 10% normal serum with 1% BSA in TBS for 2 hours at room temperature. Primary antibodies to BDNF and c-fos were applied and incubated overnight in a humidity chamber at 4°C, and then detected with rabbit anti-Mouse Alexa Fluor 488 secondary antibody for 1 hour room temperature. For DC markers, Iba1, *E.coli* and *Salmonella Thyphimurium* immunofluorescence staining, brain sections were allowed to thaw at room temperature for 10 min then fixed in cold acetone for 10 min. Tissue permeabilization was performed on all slides with PBS-0.1% Triton X-100 for 10 min at room temperature then tissues were incubated in Sudan Black B for 30 min at room temperature followed by blocking medium with PBS containing 5% BSA for 30 min at room temperature. Brain sections were incubated overnight in a humidity chamber at 4°C with rat anti-CD11b Alexa Fluor 488 antibody, Armenian hamster anti-CD103 Alexa Fluor 594 antibody and Armenian hamster anti-CD11c Alexa Fluor 647 antibody or Rabbit anti-*E.coli* antibody, Rabbit anti-*Salmonella Thyphimurium* antibody or Rabbit anti-Iba1 antibody and diluted in PBS with 1% BSA. Rabbit anti-*E.coli*, anti-*Salmonella Thyphimurium* and anti-Iba1 antibodies were then detected with donkey anti-Rabbit Alexa Fluor 488 secondary antibody or donkey anti-Rabbit Alexa Fluor 555 secondary antibody for 1 hour room temperature. For intestinal DC markers immunofluorescence staining, intestines sections were allowed to dry overnight at room temperature then fixed in 4% PFA for 10 min at room temperature. Jejunum tissues were then incubated in blocking solutions using PBS containing 5% FBS for 30 min at room temperature followed by PBS containing 10% rat serum for 30 min at room temperature. The slides were incubated overnight in a humidity chamber at 4°C with rat anti-CD11b antibody conjugated to Alexa Fluor 488 using the Mix-n-Stain^™^ CF^™^ 488A Antibody

Labeling Kit (cat no. MX488AS50, Sigma-Aldrich, Germany), rat anti-CD103 antibody conjugated to Alexa Fluor 555 using the Mix-n-Stain^™^ CF^™^ 555 Antibody Labeling Kit (cat no. MX555S20, Sigma-Aldrich, Germany) and Armenian hamster anti-CD11c Alexa Fluor 647 antibody diluted in PBS with 1% BSA. For co-staining of CD103 and *E. Coli* or *Salmonella Thyphimurium,* intestines sections were allowed to dry 10 min at room temperature then fixed in 10% Formalin for 10 min at room temperature. Jejunum tissues were then incubated in blocking solutions using PBS containing 5% FBS for 30 min at room temperature followed by protein block buffer (Agilent Dako, ON, Canada) for 30 min at room temperature. The slides were incubated overnight in a humidity chamber at 4°C with rat anti-CD103 antibody and Rabbit anti-*E.coli* antibody, or anti-*Salmonella Thyphimurium* antibody diluted in PBS with 1% BSA. Rat anti-CD103 was detected with donkey anti-Rat Alexa Fluor 488 secondary antibody, and rabbit anti-*E.coli* and anti-*Salmonella Thyphimurium* were then detected with donkey anti-Rabbit Alexa Fluor 555 secondary antibody for 1 hour room temperature. Slides were mounted in ProLong Gold with 4′,6-diamidino-2-phenylindole (DAPI) (ProLong Gold antifade regent with DAPI, Thermo Fisher Scientific, ON, Canada). Pictures were taken using an epifluorescence microscope (Eclipse 80, Nikon, ON, Canada) with the same setting and exposure time for all pictures. Confocal images were taken by Nikon A1R inverted microscope using 20x objective and 60x oil immersion objective (Nikon, ON, Canada) to detect CD103 and bacteria colocalization. Z-series image (1 μm interval, 5 – 7 μm thickness, 4096 x 4096 pixels) were obtained from jejunum and brain using 60x oil immersion objective. Quantification was performed with ImageJ software (NIH, Bethesda, MD) by counting BDNF, c-fos, CD103, CD11b or CD11c immunoreactive (positive) area Table 1.

**Table 1.**
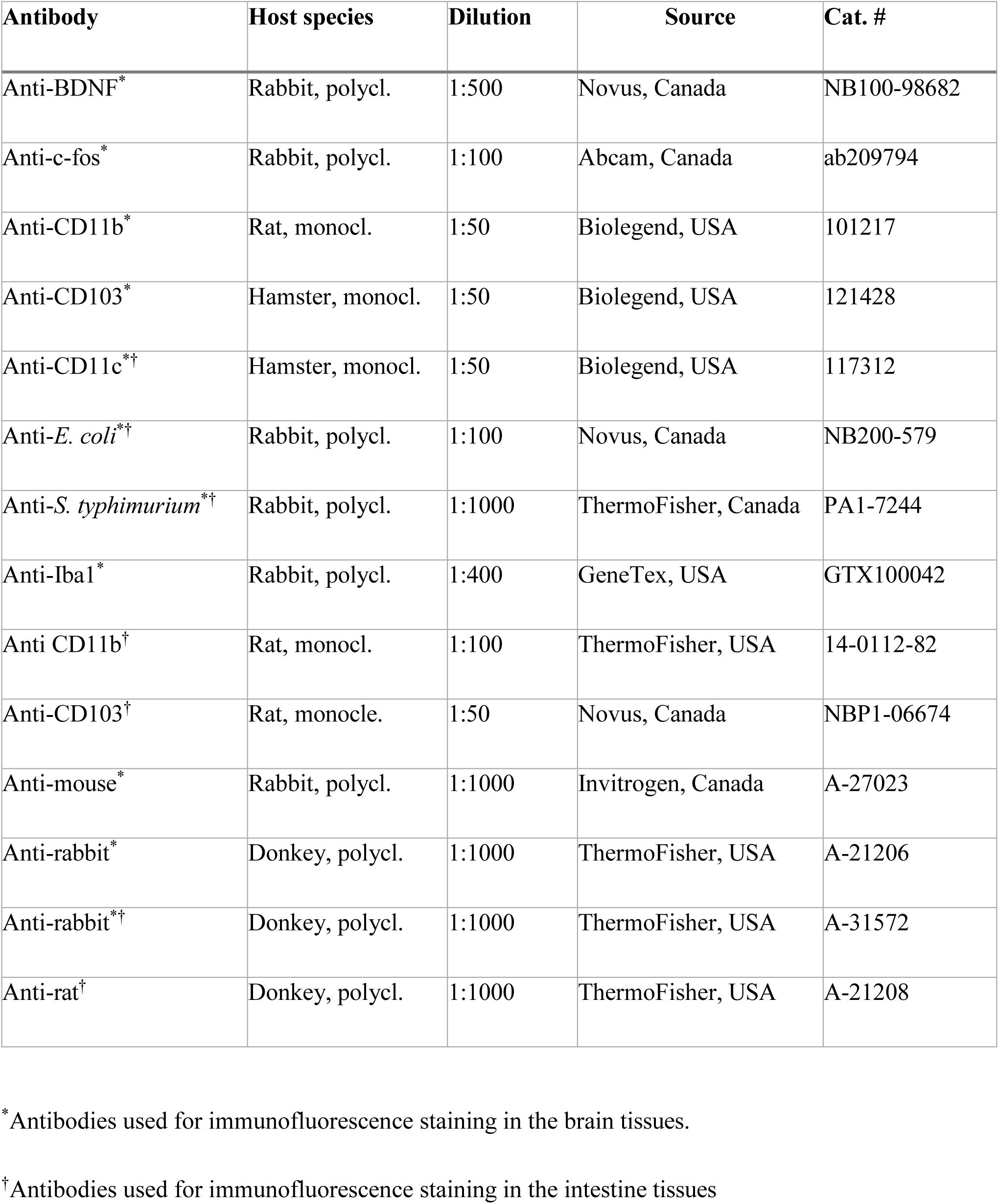
Table of antibodies used in the study.

| Antibody | Host species | Dilution | Source | Cat. # |
| --- | --- | --- | --- | --- |
| Anti-BDNF<sup>*</sup> | Rabbit, polycl. | 1:500 | Novus, Canada | NB100-98682 |
| Anti-c-fos<sup>*</sup> | Rabbit, polycl. | 1:100 | Abcam, Canada | ab209794 |
| Anti-CD11b<sup>*</sup> | Rat, monocl. | 1:50 | Biolegend, USA | 101217 |
| Anti-CD103<sup>*</sup> | Hamster, monocl. | 1:50 | Biolegend, USA | 121428 |
| Anti-CD11c<sup>*†</sup> | Hamster, monocl. | 1:50 | Biolegend, USA | 117312 |
| Anti-*E. coli*<sup>*†</sup> | Rabbit, polycl. | 1:100 | Novus, Canada | NB200-579 |
| Anti-*S. typhimurium*<sup>*†</sup> | Rabbit, polycl. | 1:1000 | ThermoFisher, Canada | PA1-7244 |
| Anti-Iba1<sup>*</sup> | Rabbit, polycl. | 1:400 | GeneTex, USA | GTX100042 |
| Anti CD11b<sup>†</sup> | Rat, monocl. | 1:100 | ThermoFisher, USA | 14-0112-82 |
| Anti-CD103<sup>†</sup> | Rat, monocle. | 1:50 | Novus, Canada | NBP1-06674 |
| Anti-mouse<sup>*</sup> | Rabbit, polycl. | 1:1000 | Invitrogen, Canada | A-27023 |
| Anti-rabbit<sup>*</sup> | Donkey, polycl. | 1:1000 | ThermoFisher, USA | A-21206 |
| Anti-rabbit<sup>*†</sup> | Donkey, polycl. | 1:1000 | ThermoFisher, USA | A-31572 |
| Anti-rat<sup>†</sup> | Donkey, polycl. | 1:1000 | ThermoFisher, USA | A-21208 |

\* Antibodies used for immunofluorescence staining in the brain tissues.
† Antibodies used for immunofluorescence staining in the intestine tissues

### Gene expression study

RNA from colon and brain (hippocampus and amygdala) tissues were extracted using the RNeasy Mini Kit (Qiagen, Toronto, Canada) and DNase digestion during purification was carried out using the RNase-free DNase (Qiagen). NanoString nCounter® Gene Expression CodeSet for mouse inflammation genes and a custom Nanostring gene expression codeset for selected genes were run on colon and brain tissues, respectively, according to manufacturer’s instructions (NanoString Technologies, Inc., Seattle, WA). The results obtained were analyzed with the analysis software nSolver 2.5 (NanoString Technologies). The Log2 ratios built from the data obtained were then uploaded into Ingenuity Pathway analysis software (Qiagen) for further analysis. The network score is based on the hypergeometric distribution and is calculated with the right-tailed Fisher’s exact test.

### Label-Free Quantitative Proteomics Assay using Nano-UPLC MS

Around 10 mg of brain tissue was homogenized by sonication in 150 μL of lysis buffer with 50 mM ammonium bicarbonate, pH 8, 0.25% Rapigest SF (Waters), complete protease inhibitor (Roche), and phosphatase inhibitor cocktails I and III (Sigma-Aldrich). After centrifuge, the supernatants were collected and protein concentrations were assayed by Bradford assay. To quantitatively determine proteins expression for proteomic analysis, each sample was diluted to 5 mg/mL with lysis buffer and was reduced by incubation with 10 mM DTT for 10 min at 80 °C. Then iodoacetamide was added to 20 mM, and samples were incubated at room temperature in the dark for 30 min. Sequencing-grade modified trypsin (Promega) of 1:100 (w/w) was added to digest proteins samples under incubation at 37 °C for overnight. Next, samples were acidified to a final concentration of 1% TFA / 2% ACN and continually incubated at 60 °C for another 2 h. Samples were centrifuged at 20, 000g for 10 min to remove any undissolved residue. Finally, 50 μg of digested proteins per sample were transferred to Total Recovery LC Vials (Waters), and 50 fmol of MassPrep ADH standard (Waters) per μg of brain protein digests were added as an internal standard. Digested peptides were purified and analyzed using reverse phase uplc (nanoacquity) with qtof ms/ms (Waters) as previously described ^28^.

### Statistical analysis

Statistical analysis was performed using Prism 6 software. Statistical comparisons were performed using one-way ANOVA, student t-test, Wilcoxon’s signed rank t-test or multiple t-test as appropriate. One-way ANOVA followed by Dunn’s test for multiple comparison was used to compare GF, colonized (*E. coli*, ASF, SPF) mice, and pcSPF. Immunofluorescence was analyzed using Mann-Whitney t-test or One-way ANOVA followed by Dunn’s test for multiple comparisons. Behavior before and after LPS and poly I:C, cosalane, fingolimod or resatorvid treatment were compared using Wilcoxon’s signed rank t-test. Benjamini and Hochberg FDR correction method was used when multiple comparisons were performed for NanoString. For all box-plot graphs, the median is represented by the center line with 10 - 90 percentile whiskers. A p-value of <0.05 was considered statistically significant.

## Supplementary Figures

**Supplementary Fig. 1.**
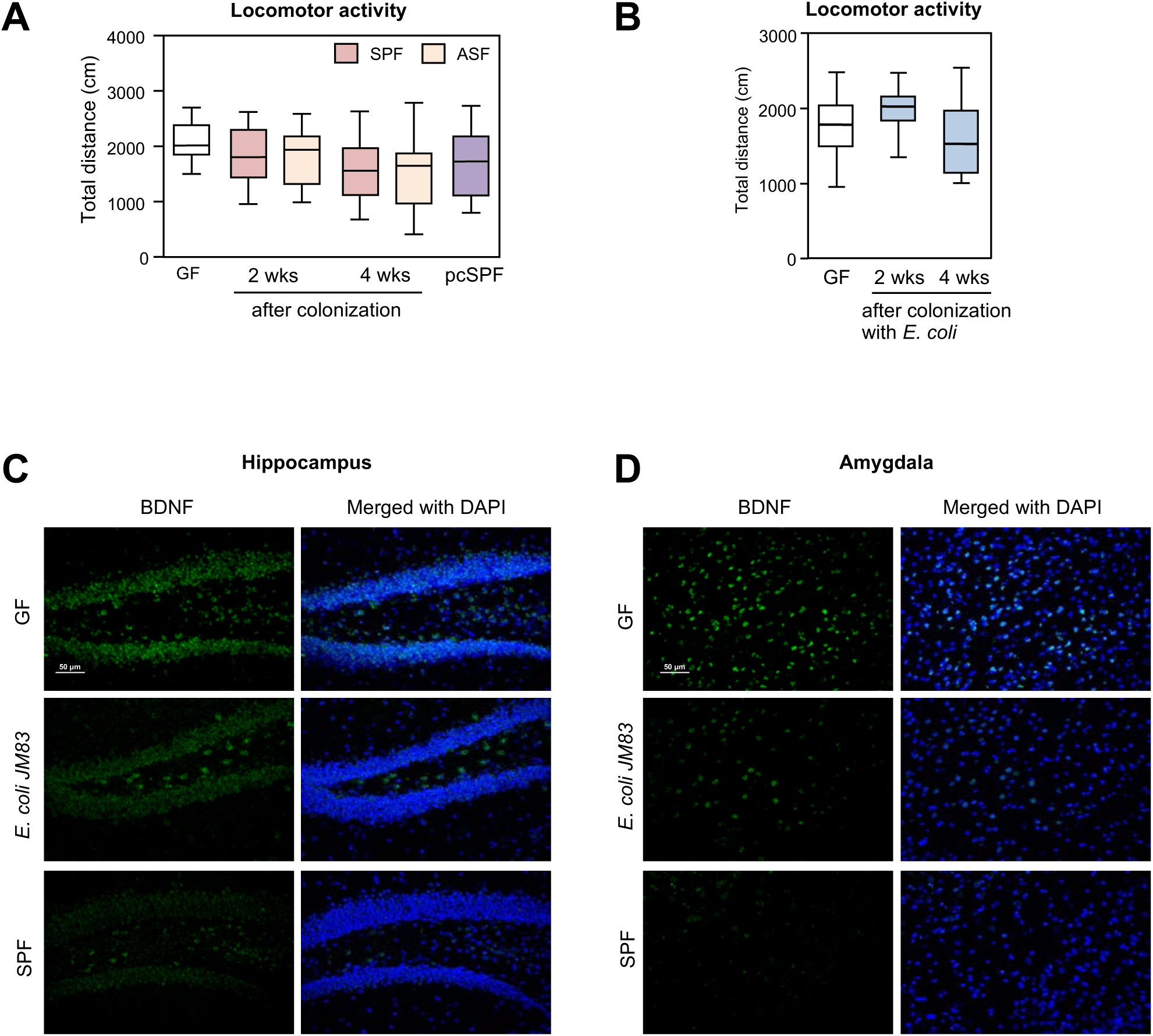
Locomotor activity in mice colonized with SPF and ASF microbiota and E. coli JM83 and effect of the mono-colonization on BDNF expression in the brain. A) Locomotor activity of GF mice colonized with ASF and SPF microbiota and of conventionally raised mice (pcSPF) in the light/dark preference test (n=16/group). B) Locomotor activity in the light/dark preference test of GF mice mono-colonized with *E. coli* JM83 (n=18). C) Representative images of the hippocampal BDNF expression in GF, E. coli mono-colonized and SPF mice. D) Representative images of the amygdala BDNF expression in GF, E. coli mono-colonized and SPF mice.

**Supplementary Fig. 2.**
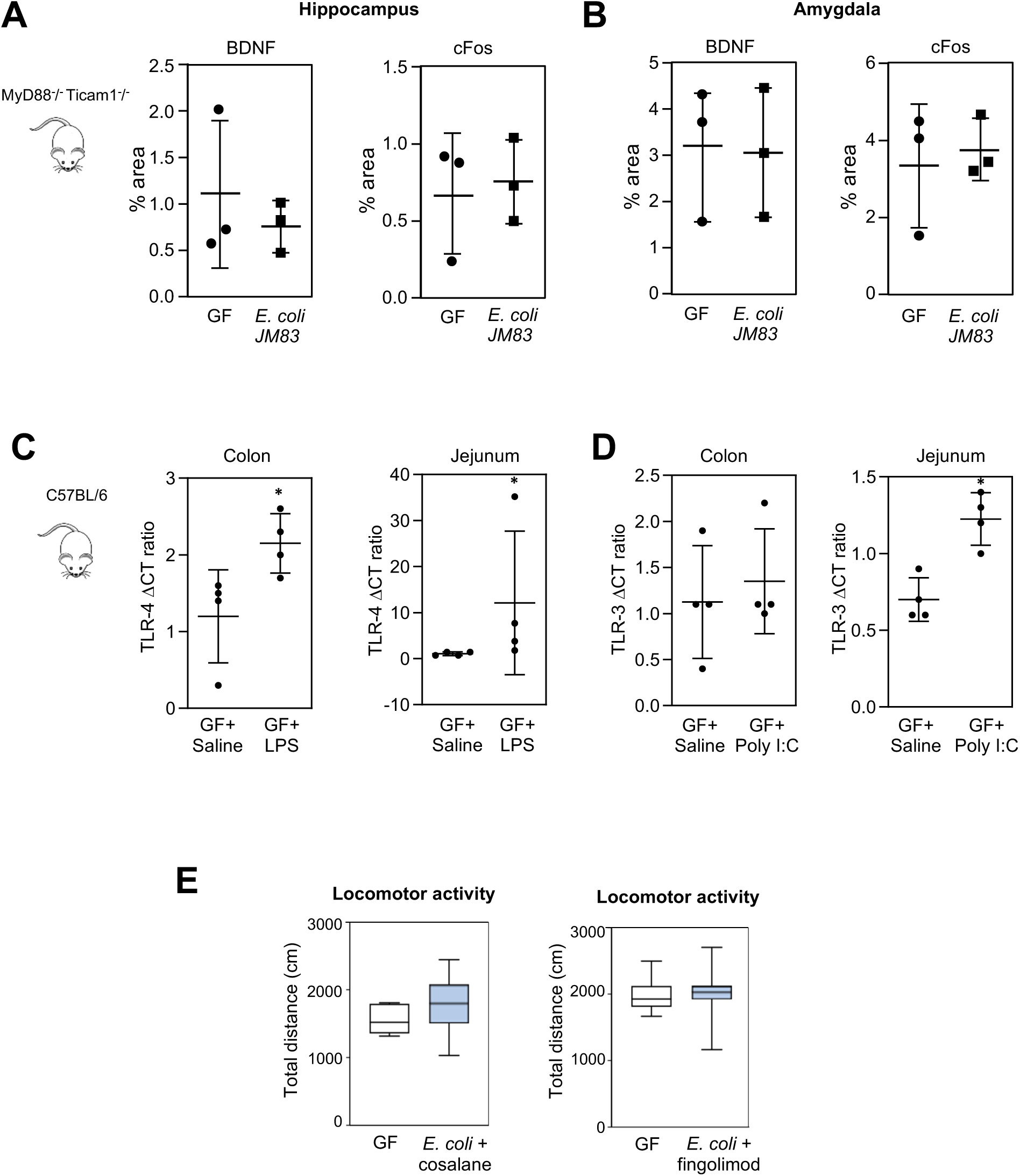
Brain BDNF and c-fos expression in mono-colonized MyD88^−/−^ Ticam1^−/−^ mice, TLR4 and TLR3 gene expression in LPS and Poly I:C treated mice and locomotor activity in E. coli mono-colonized treated with cosalane and fingolimod. A) Quantification of BDNF and c-fos expression in hippocampus and amygdala (B) of mono-colonized MyD88^−/−^ Ticam1^−/−^ mice. C) Quantification of TLR4 gene expression in colon and jejunum of LPS treated mice. D) Quantification of TLR3 gene expression in colon and jejunum in Poly I:C treated mice. E) Locomotor activity (total distance travelled) of GF mice mono-colonized with *E. coli* JM83 treated with cosalane or fingolimod (n=7/group). Statistics by Wilcoxon’s signed rank test or t-test or Mann-Whitney test, *p<0.05 vs GF.

**Supplementary Fig. 3.**
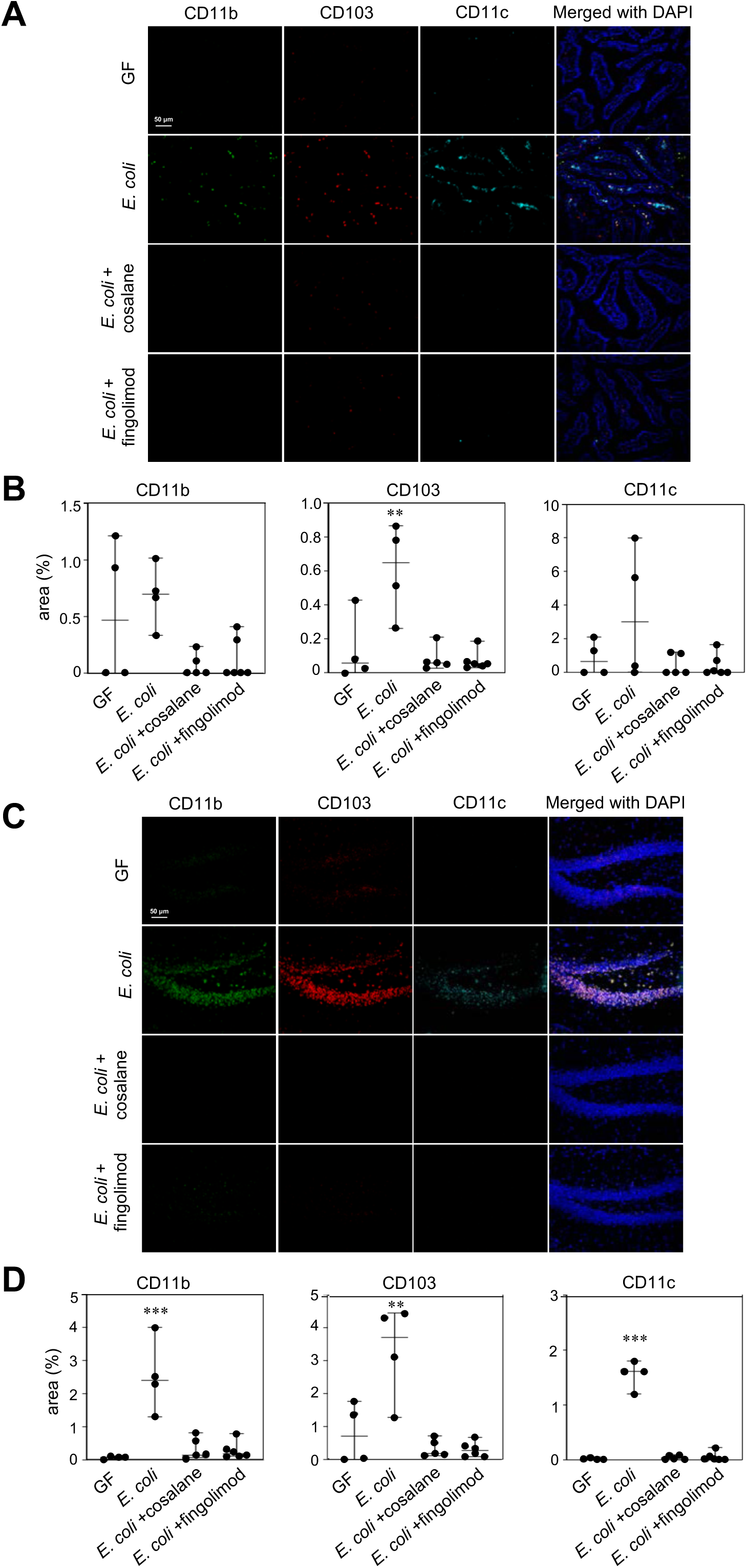
Effects of cosalane and fingolimod treatment on CD11b, CD103 and CD11c expression in the hippocampus and jejunum. A, B) CD11b, CD103 and CD11c expression using immunofluorescence in the jejunum lamina propria of GF (n=4), *E. coli* JM83 mono-colonized mice treated with saline (n=4), cosalane (n=5) or fingolimod (n=6). C, D) CD11b, CD103 and CD11c expression using immunofluorescence in the hippocampus (dentate gyrus) of GF (n=4) and E. coli JM83 mono colonized mice treated with saline (n=4), cosalane (n=5) or fingolimod (n=6). Statistics by One-way ANOVA, **p<0.01, ***p<0.001 vs GF, cosalane and fingolimod groups.

**Supplementary Fig. 4.**
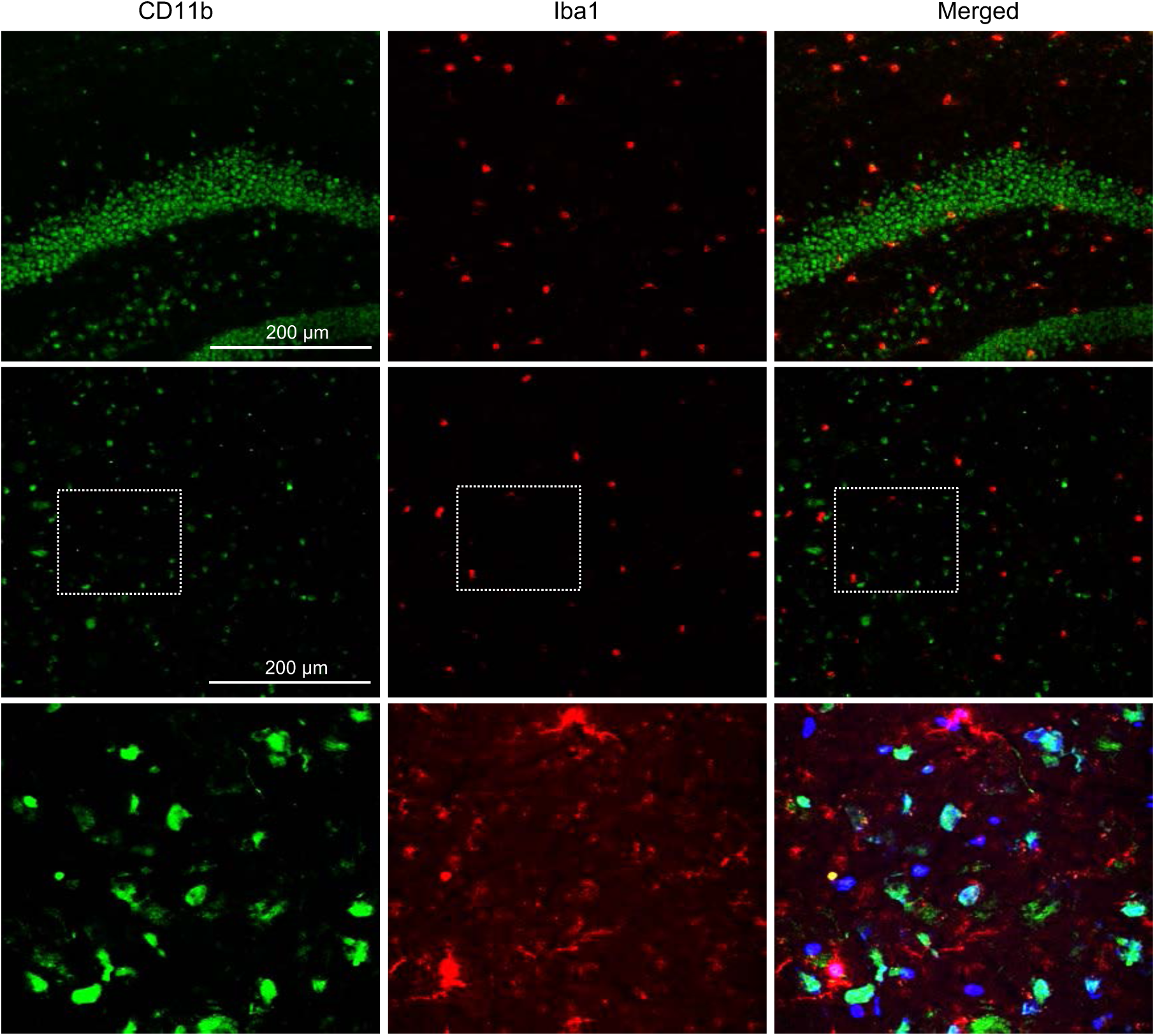
Immunofluorescence staining for dendritic cells and microglia. Representative images of immunofluorescent staining of CD11b+ cells (green) and Iba1+ cells (red) in the hippocampus (upper panel) or amygdala (middle panel) of a GF mouse mono-colonized with *E. coli* JM83. The lower panel shows higher magnification (x60) images of the amygdala. White squares in the middle panel indicate the magnified areas displayed in the lower panel.

**Supplementary Fig. 5.**
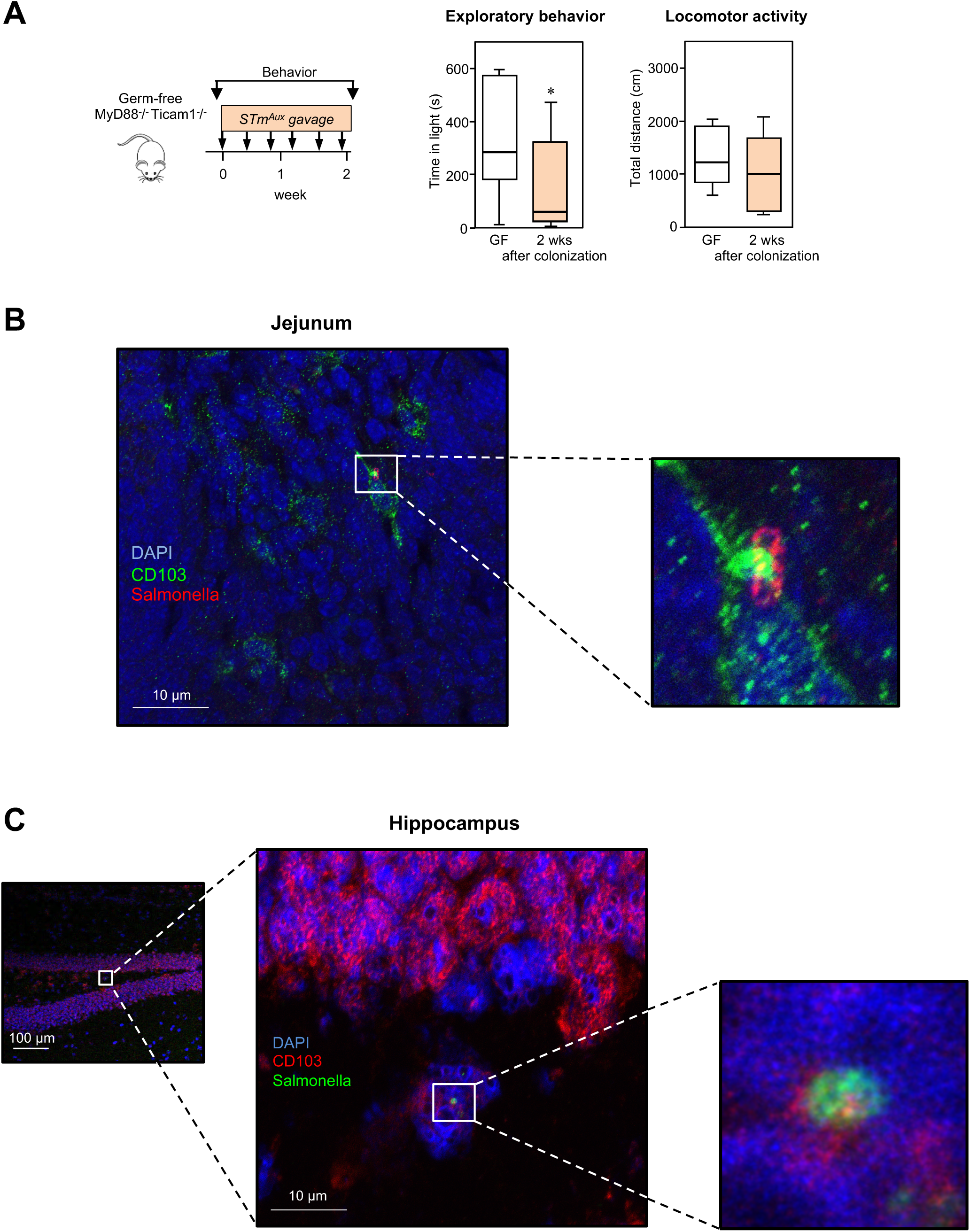
Behavior of GF MyD88^−/−^ Ticam1^−/−^ mice mono-colonized with transient Salmonella enterica serovar Typhimurium and bacteria presence in the intestine and the brain. A) Behavior of GF C57BL6 MyD88^−/−^ Ticam1^−/−^ mice before and after colonization with auxotrophic *Salmonella enterica* serovar *Typhimurium* (StmAux). Statistics by Wilcoxon’s signed rank test; *p<0.05 vs GF; n=8. B) Representative images of immunofluorescent staining of CD103+ cells (green) and Salmonella+ cells (red) in the jejunum of GF mice mono-colonized with StmAux. White squares indicate the magnified areas. C) Representative images of immunofluorescent staining of CD103+ cells (red) and Salmonella+ cells (green) in the hippocampus of GF mice mono-colonized with StmAux. White squares indicate the magnified areas.

**Supplementary Fig. 6.**
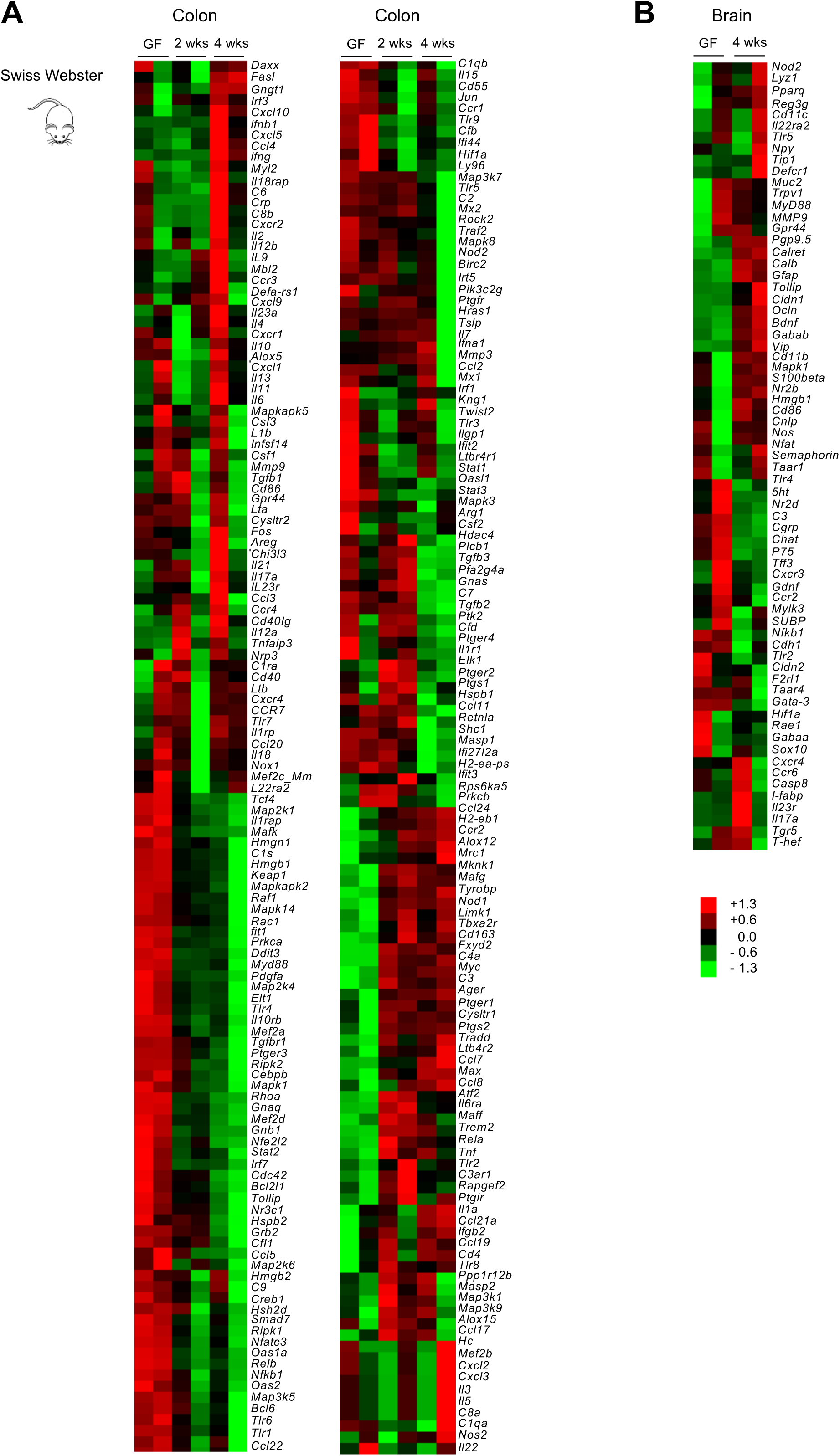
Multiple innate immunity and neural system related genes are altered in the colon and brain of Swiss Webster mice following bacterial colonization. A) 250 inflammatory genes from colon tissues of Swiss Webster GF and E. coli HA107 mono-colonized mice using the NanoString nCounter® CodeSet for mouse inflammation. B) 72 neuro-immuno-epithelial genes from brain tissues of Swiss Webster GF and *E. coli* HA107 mono-colonized mice using a customized NanoString nCounter® CodeSet.

**Supplementary Fig. 7.**
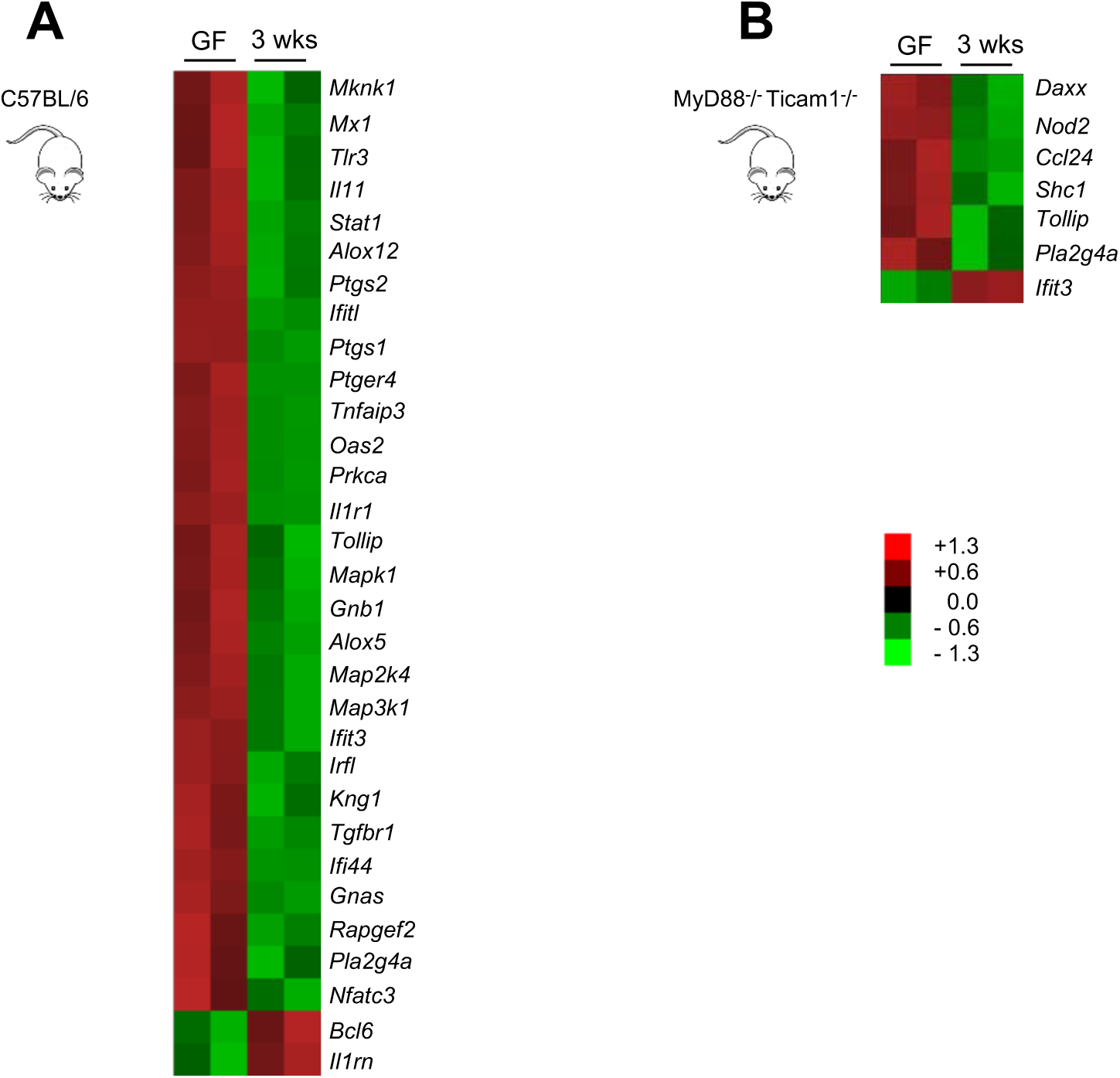
Multiple immunity related genes significantly altered in colon of C57BL/6 mice and MyD88^−/−^ Ticam1^−/−^ mice following bacterial colonization. A) Inflammatory genes significantly different between colon tissues of GF C57BL/6 and C57BL/6 *E. coli* mono-colonized mice using the NanoString nCounter® CodeSet for mouse inflammation. B) Inflammatory genes significantly different between colon tissues of GF MyD88^−/−^ Ticam1^−/−^ and MyD88^−/−^ Ticam1^−/−^ *E. coli* mono-colonized mice using the NanoString nCounter® CodeSet for mouse inflammation.

**Supplementary Fig. 8.**
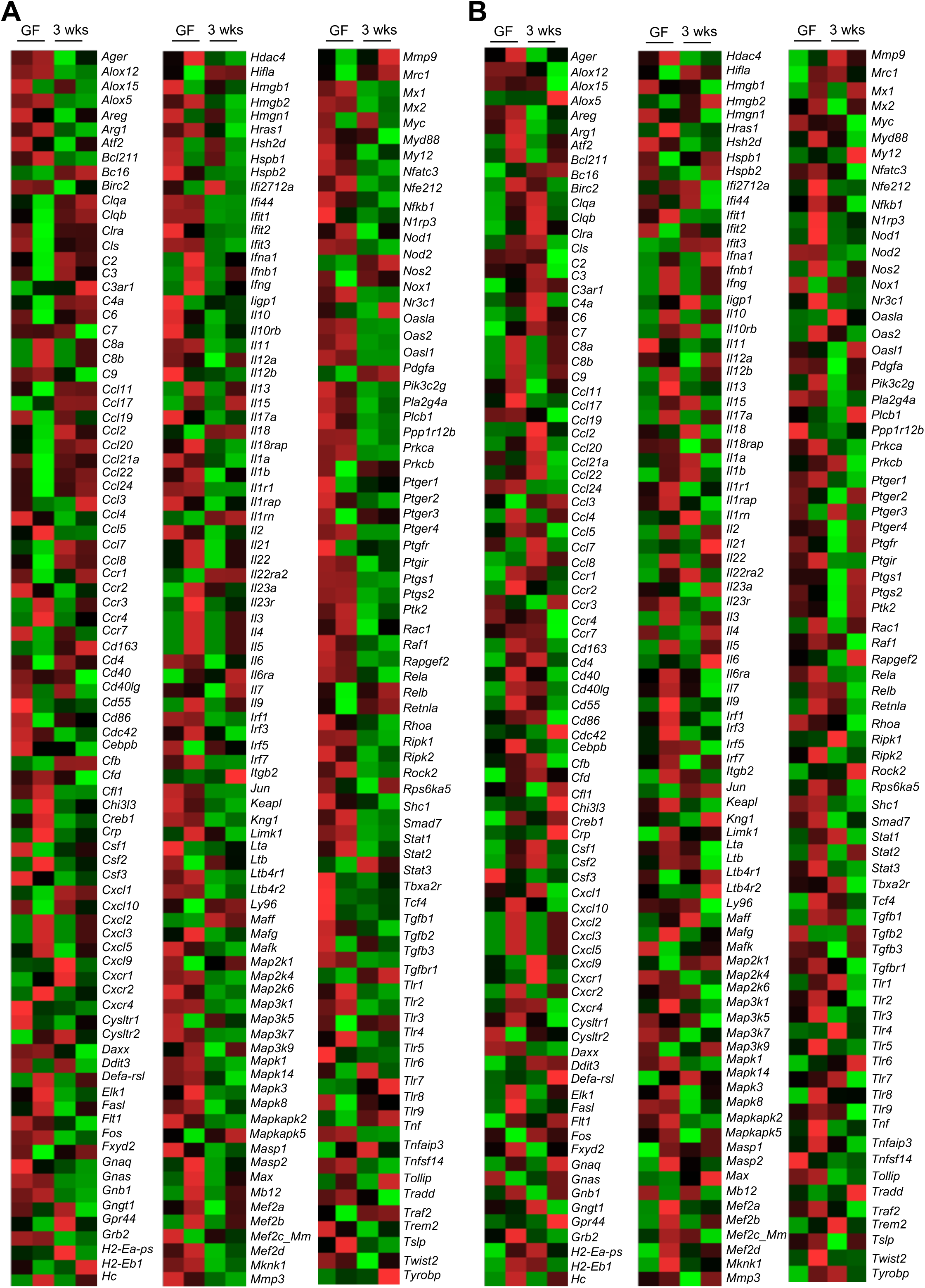
Immunity related genes expression in colon of C57BL/6 mice and MyD88^−/−^ Ticam1^−/−^ mice following bacterial colonization. A) Colonic inflammatory genes from GF C57BL/6 and C57BL/6 *E. coli* mono-colonized mice using the NanoString nCounter® CodeSet for mouse inflammation. B) Colonic inflammatory genes from GF MyD88^−/−^ Ticam1^−/−^ and MyD88^−/−^ Ticam1^−/−^ *E. coli* mono-colonized mice using the NanoString nCounter® CodeSet for mouse inflammation.

**Supplementary Fig. 9.**
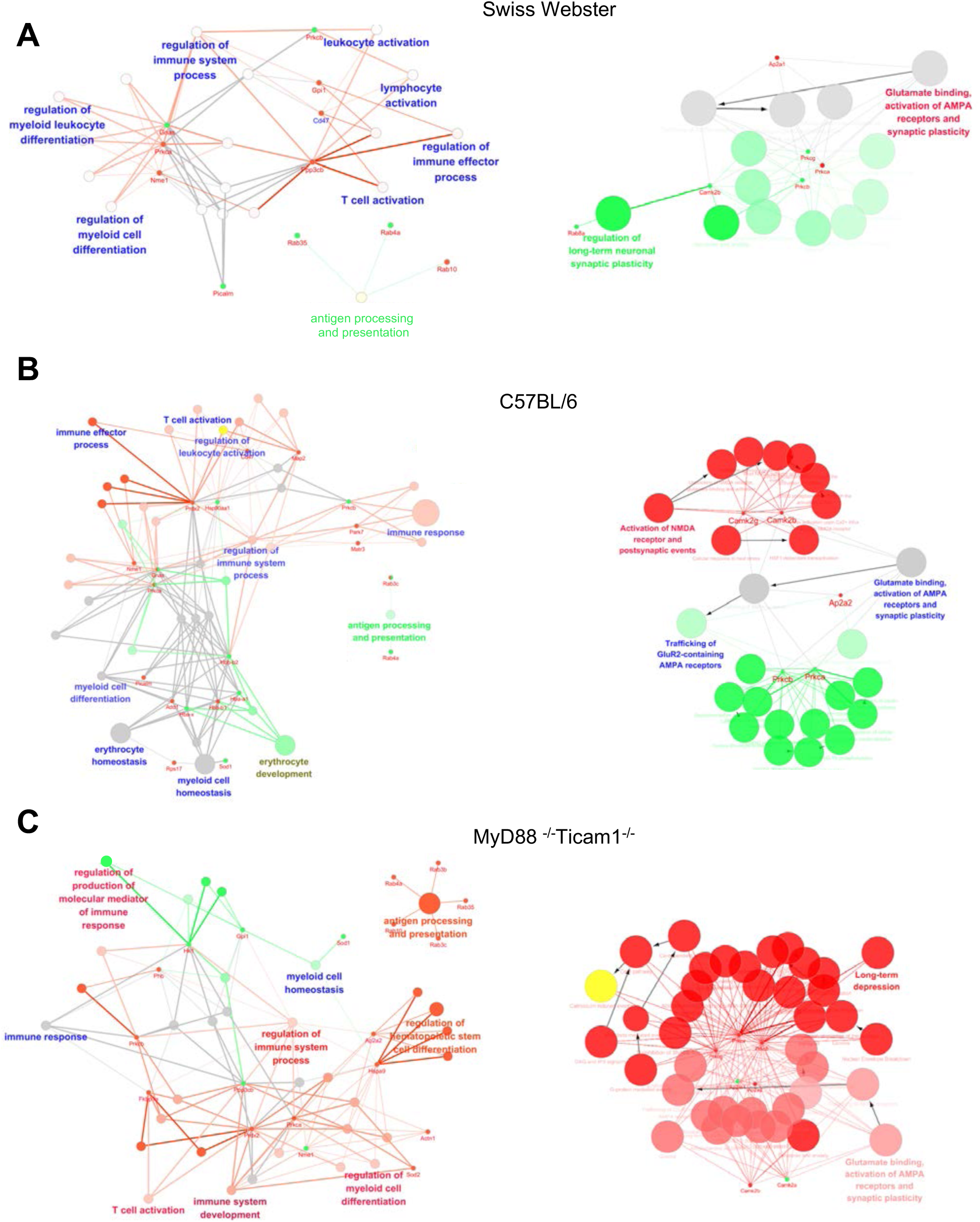
Changes in brain immune and neural networks following bacterial monocolonization assessed by proteomics. Schematic illustration of systematic rewiring of dynamic biological networks by analyzing differently expressed proteins following bacterial monocolonization with *E. coli* HA107 in Swiss Webster mice (A), C57BL/6 mice (B) and MyD88^−/−^ Ticam1^−/−^ mice (C) related to immune system function (left) and neural plasticity (right) (n=3/group). Network analysis was performed with ClueGo and visualized with Cytoscape. The mean expression level for each protein is visualized with small dot (green for increased, red for decreased expression). Biological pathways and functions are visualized as colored circular nodes linked with related proteins based on their change of expression level, up-regulated shown in green while those down-regulated in red.

**Supplementary Table 1.**
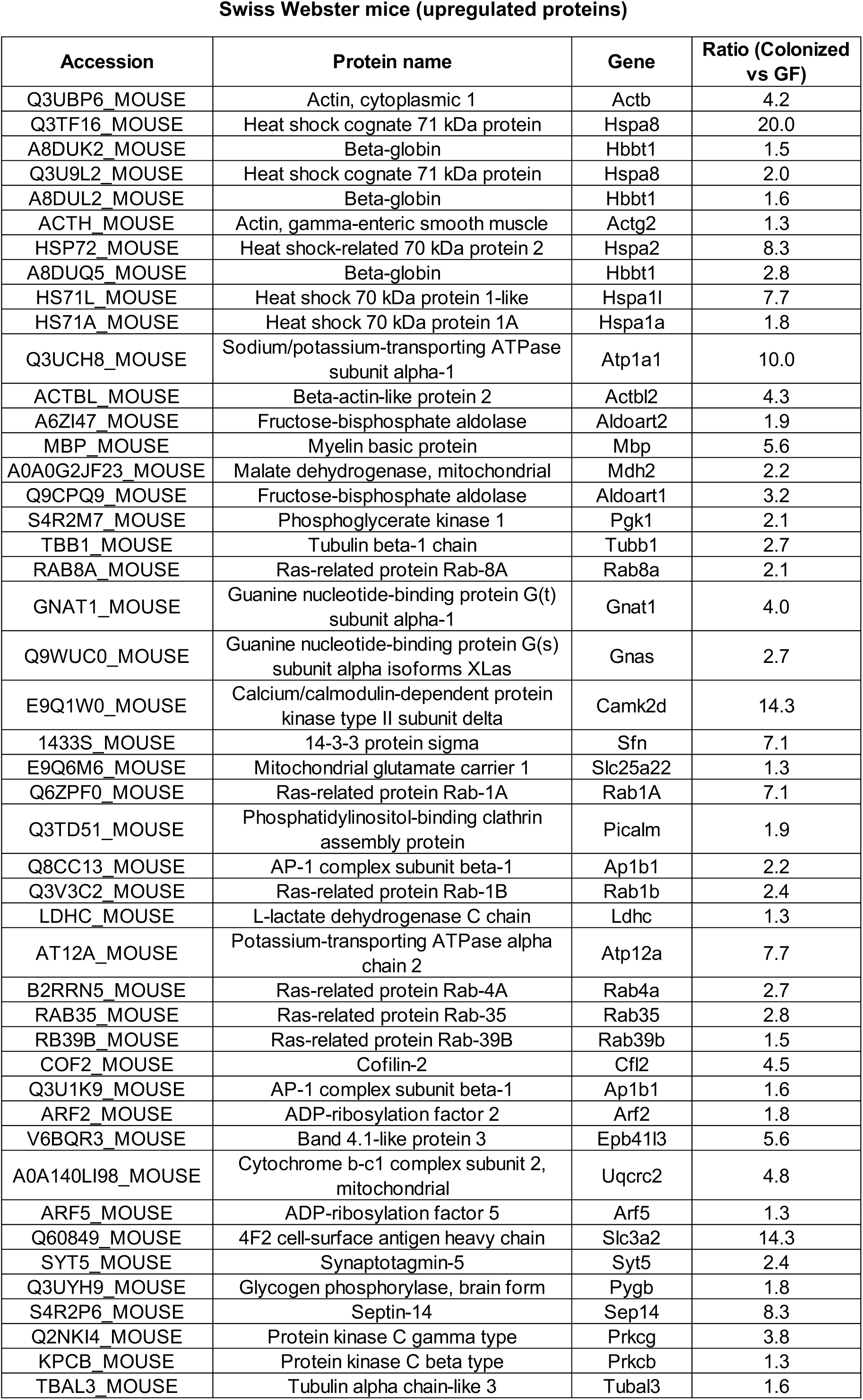
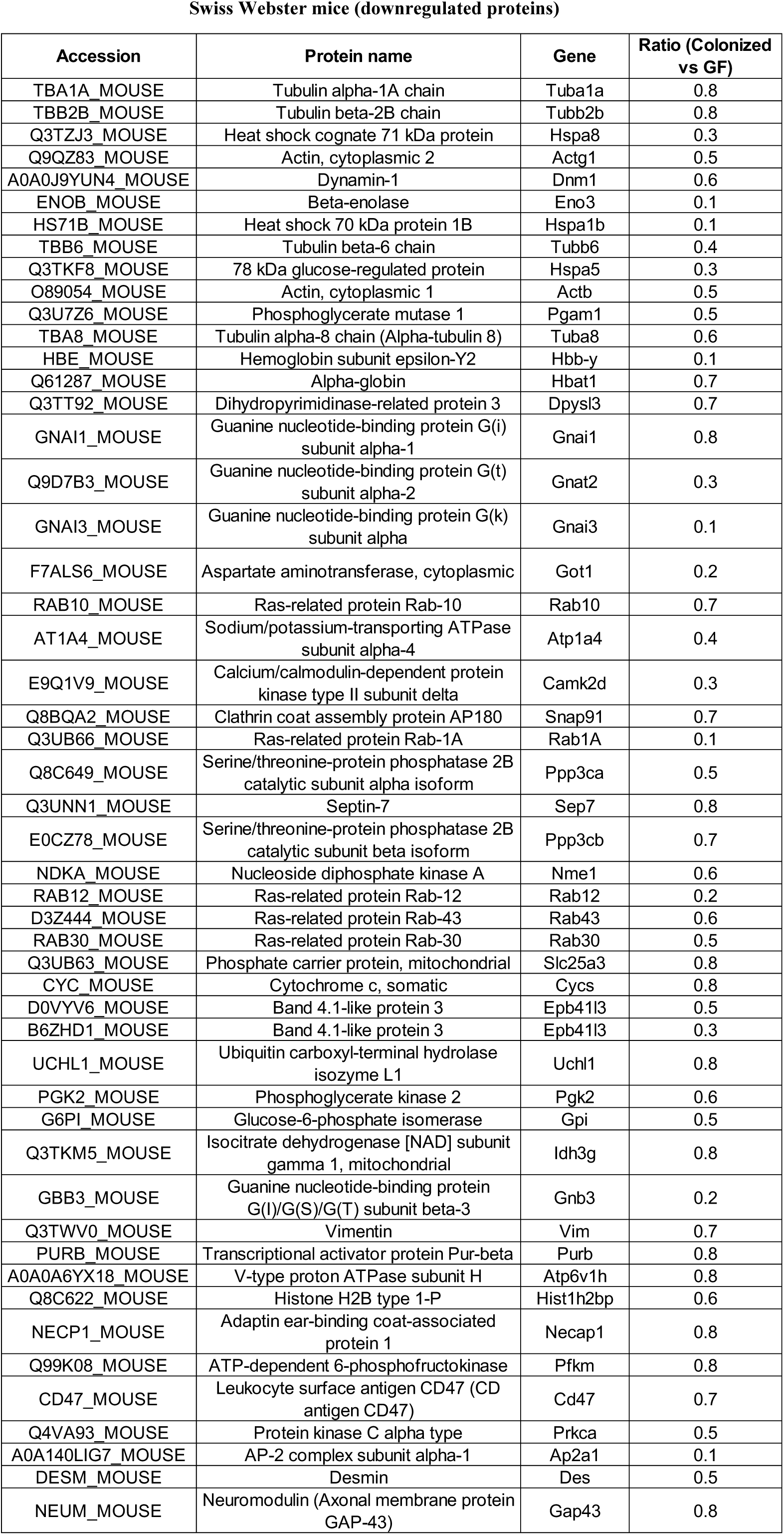
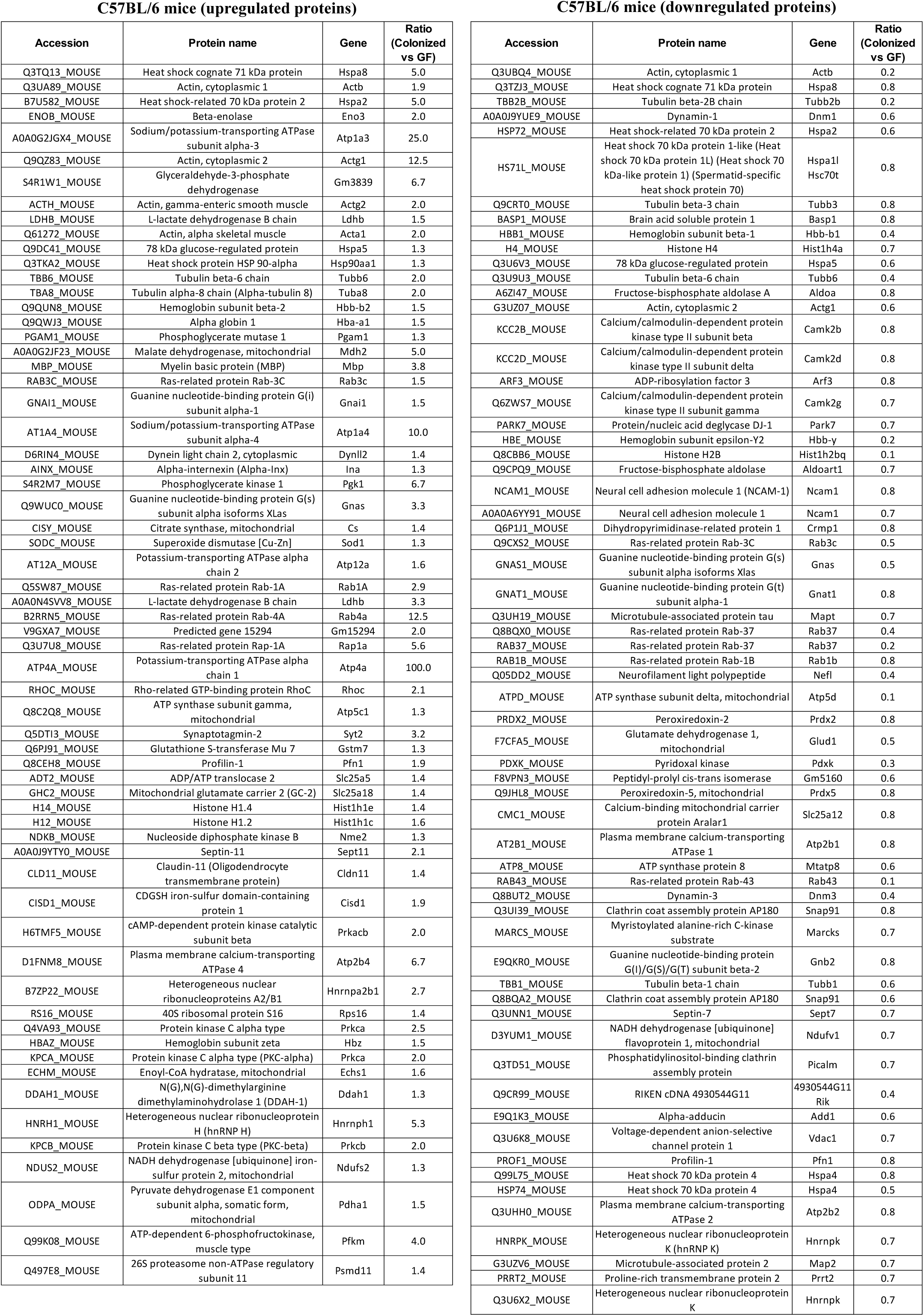
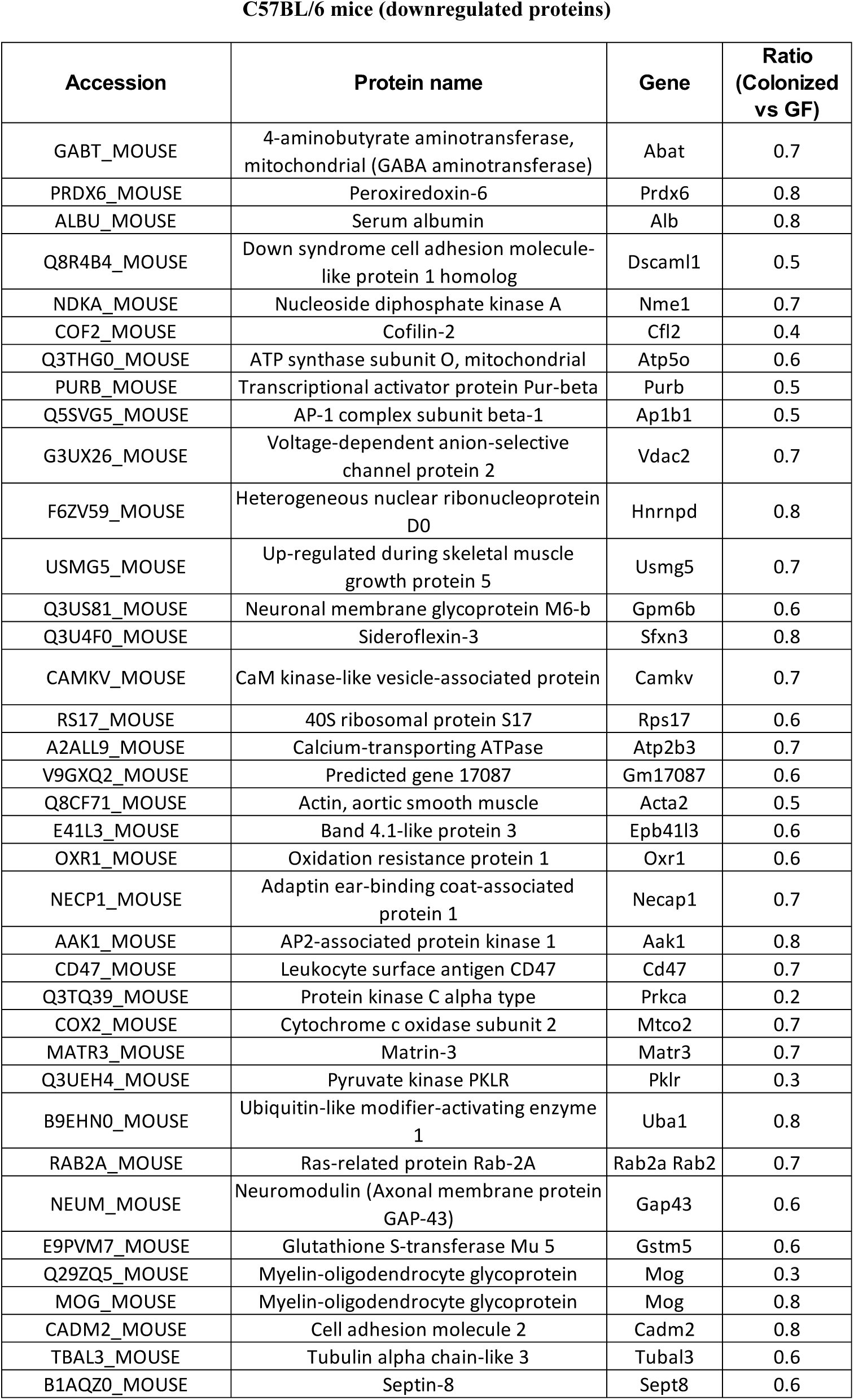
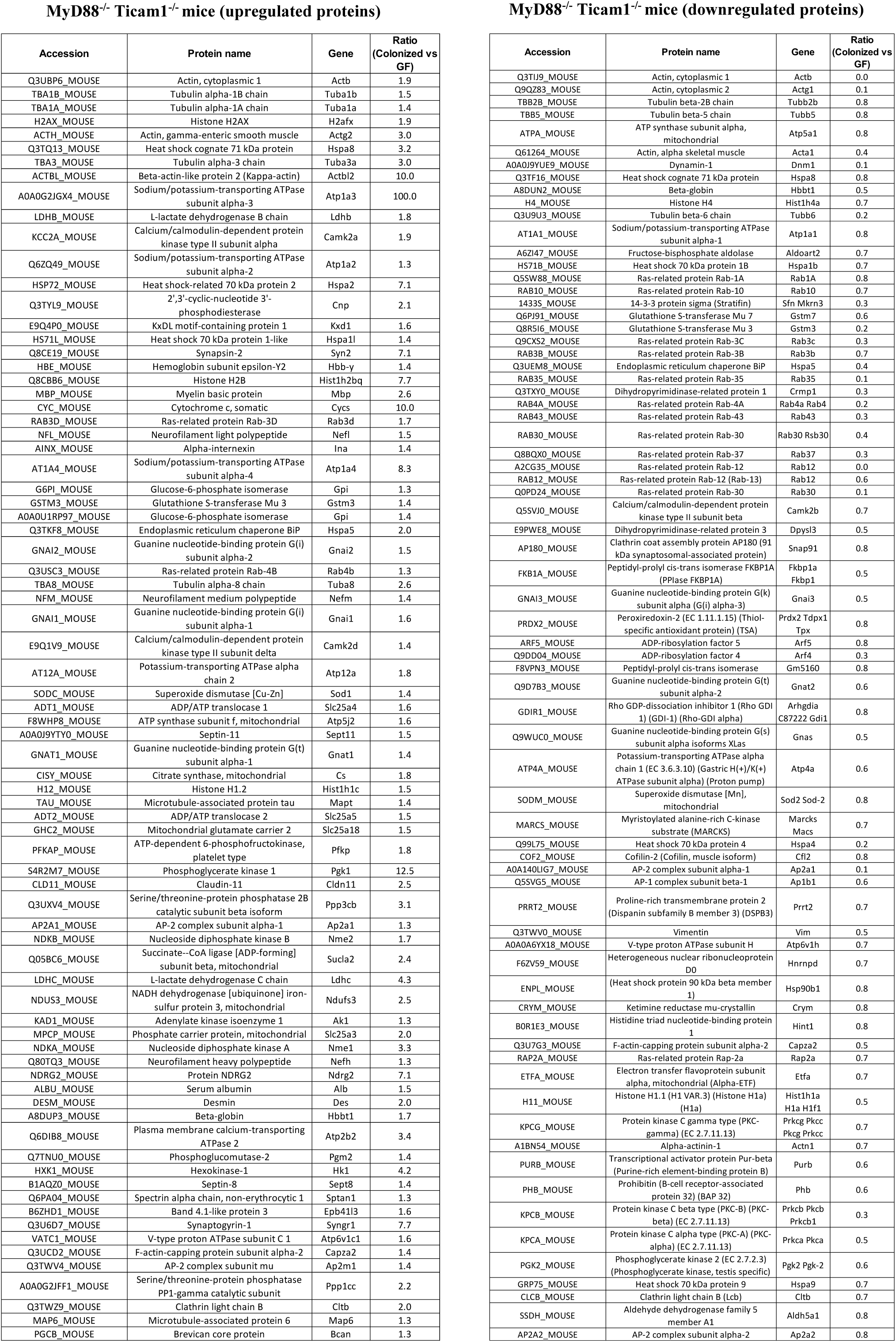
List of proteins that are significantly altered compared to GF mice. Swiss Webster mice: 492 proteins were identified of which 97 proteins significantly altered compared to GF; 51 downregulated and 46 upregulated. C57BL/6 mice: 462 proteins were identified of which 163 proteins significantly altered compared to GF; 100 downregulated and 63 upregulated. MyD88−/− Ticam1−/− mice: 439 proteins were identified of which 149 proteins significantly altered compared to GF; 72 down-regulated and 77 up-regulated. Ratio >1 is upregulated proteins; ratio<1 is downregulated proteins compared to GF (n=3/group), (p<0.05 vs GF).

## Notes

### Competing Interest Statement

The authors have declared no competing interest.

